# An injury-responsive *bHLH* is required for regenerative responses following mechanical injury in adult *Schistosoma mansoni*

**DOI:** 10.64898/2026.07.31.742058

**Authors:** Lu Zhao, James. J. Collins

**Author notes:** To whom correspondence should be addressed James J. Collins III UT Southwestern Medical Center Department of Pharmacology 6001 Forest Park. Rd. Dallas, TX 75390, USA.

## Abstract

Adult schistosomes can survive for decades in the hostile niche of the host vasculature, yet the mechanisms that support tissue repair and resilience remain poorly understood. Here, we show that both mechanical and chemical injury trigger robust proliferative responses in adult parasites and display distinct spatial patterns. We identify a *bHLH* transcription factor that responds specifically to mechanical injury and is required for injury-induced stem cell proliferation and parenchymal cell renewal. Downstream, we find a WD40 repeat-containing protein that acts within a subset of stem cells to mediate this regenerative program. Together these findings detail an injury-responsive regenerative program in schistosomes that supports both their longevity and resilience *in vivo*.

## Introduction

Schistosome infections affect more than 250 million people worldwide, with approximately 150 million untreated (*1*), often leading to chronic and debilitating disease (*2, 3*). A striking feature of these parasites is their remarkable longevity: adult schistosomes can persist within their host for decades (*3*), despite continuous exposure to the host immune response and mechanical and chemical stress within the bloodstream. How these parasites maintain tissue integrity in such a hostile environment remains an open question.

One explanation is that schistosomes possess limited regenerative capability. Although they lack the extensive regenerative potential seen in free-living flatworm planarians and cannot regenerate new heads or tails post amputation (*4*), studies suggest that schistosomes do possess some capacity for tissue repair following injury. Adult schistosomes can rapidly form new tegument (skin) after physical injury (*5*) and recover from sublethal praziquantel (PZQ) exposure through tissue repair *in vivo* (*6*). These observations suggest that tissue repair could be important for parasite survival, raising the possibility that regenerative-like mechanisms contribute to their long-term persistence in the blood. However, the molecular basis of these regenerative responses remains unknown.

Somatic stem cells, known as neoblasts, are required for maintaining adult schistosome tissues, most notably the tegument and the gut (*7–9*). Following mechanical injury by puncture with a sharpened needle, schistosome neoblasts undergo massive proliferation at the wound site (*10*); however, it is not clear whether injury-induced neoblasts proliferation drives tissue repair. In intact worms, neoblast activity is largely biased toward tegument maintenance (*8*). However, it is tempting to speculate that injury affecting multiple tissues may shift neoblast developmental potential to support broader regenerative responses beyond the tegument. This idea is supported by developmental juvenile stages, during which neoblasts display expanded molecular diversity, suggesting broader development potential(*11*). Notably, outside the testis, *eled*^+^ neoblasts are sparsely distributed and restricted to the gut region in adult worms (*9*), whereas they are abundant in juvenile worms, particularly in posterior regions undergoing rapid growth (*11*). This raises the possibility that these *eled*^+^ neoblasts represent a more plastic population with broader developmental potential that may enable adult worms to repair tissue damage following injury. However, whether these cells are reactivated in response to injury in adult worms, and how injury signals regulate their behavior, remain unknown.

Here we show that adult *Schistosoma mansoni* mount robust proliferative and transcriptional responses to multiple forms of injury. Mechanical injury rapidly induces a *bHLH* transcription factor whose expression shifts from tegument progenitors to mature tegument cells. Functional analyses demonstrate that *bHLH* is required for injury-induced neoblast activation and the generation of new parenchymal cells. We further identify a WD40 repeat-containing protein that acts downstream within *eled* neoblasts to mediate this regenerative program. Together, these findings define a previously unrecognized injury-responsive transcriptional program that links tissue damage to neoblast activation and provides new insight into mechanisms underlying parasite longevity *in vivo*.

## Results

### Different types of injury induce proliferative responses in adult *S. mansoni*

To test whether different types of injury could stimulate neoblast proliferation, we subjected worms to mechanical (puncture with a needle or amputation) or chemical (sublethal PZQ, 1µg/mL for 6hrs) injuries. Consistent with previous studies (*10*), we observed a localized accumulation of EdU^+^ cells at the site of puncture injuries following 2-day recovery, confirming our previous observation that physical tissue damage triggers proliferative responses (Fig. 1A). Similarly, amputation of worms with a sharpened probe likewise resulted in a marked accumulation of EdU^+^ cells adjacent to the site of injury (Fig. 1A). Instead of a localized response that accompanies puncture or amputation type injuries, we observed that sublethal PZQ exposure induces a broader proliferative response, with EdU^+^ cells distributed throughout the entire trunk of the worms (Fig. 1A).

**Figure 1.**
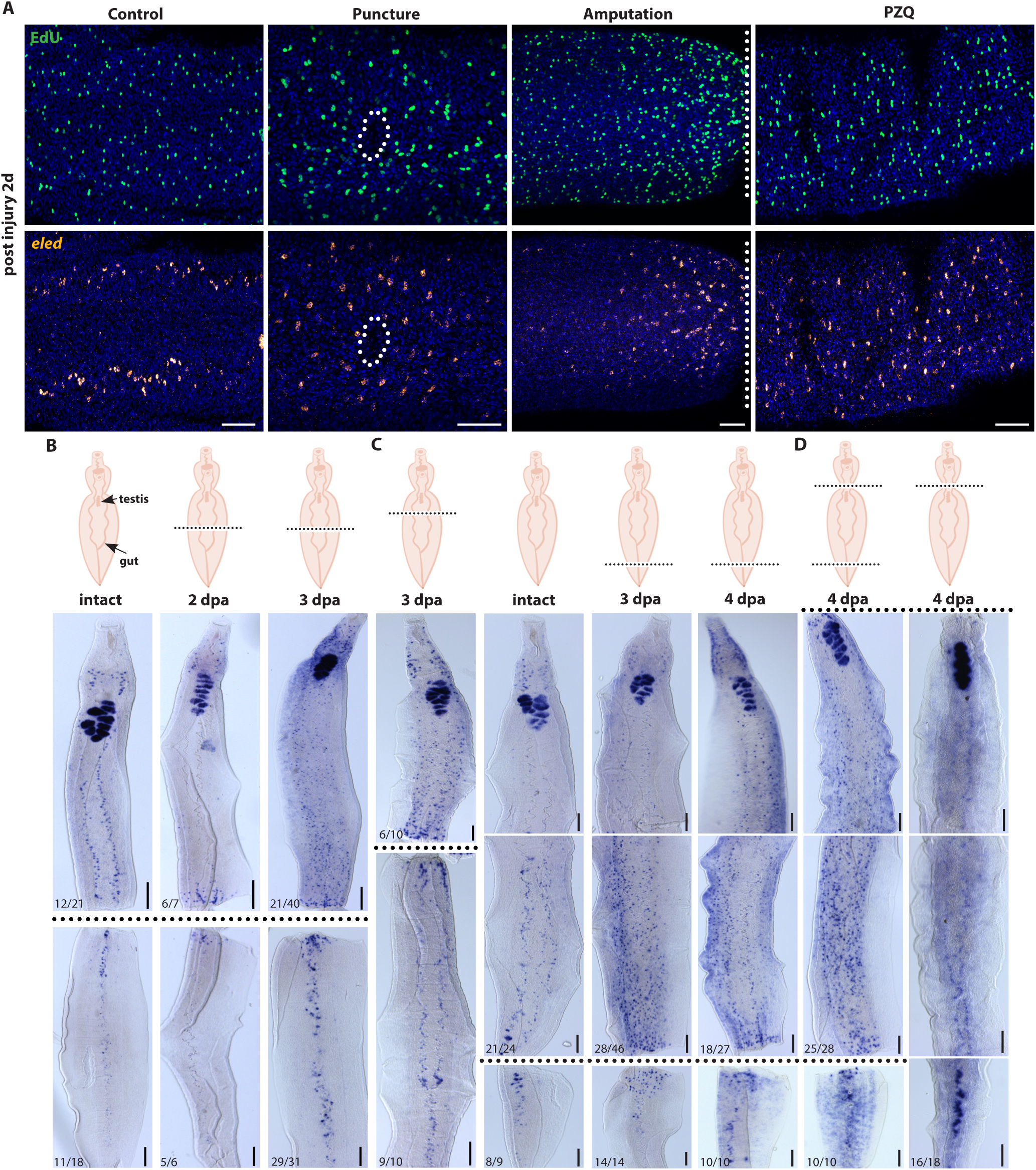
Different types of injury induce proliferative responses in adult *S. mansoni*. (**A**) Fluorescence *in situ* hybridization (FISH) showing EdU*^+^* and *eled*^+^ neoblasts following mechanical and chemical injury at 2 days post-injury in adult *S. mansoni*. Mechanical injury was induced either by needle puncture or mid-trunk amputation. Dashed line approximates the site of injury. Chemical injury was induced by exposing worms to 1µg/mL praziquantel (PZQ) for 6 hrs. Representative images are shown for puncture injury (intact: n=7/8 males, puncture: n =9/9 males), amputation (intact: n=10/10, amputation: n=11/12), PZQ exposure (EtOH: n=32/37, PZQ: n=34/47). Data were collected from at least two biological replicates. Scale bar: 50 µm. (**B-D**) Top, schematic illustrating different amputation strategies. Bottom, colorimetric whole-mount *in situ* hybridization (WISH) showing *eled* expression in mid-trunk-amputated worms at (B) 2 and 3 days post-amputation (dpa), (C) anterior-trunk-amputated and tail-amputated worms at 3 and 4 dpa, and (D) tail-and-head-amputated and head-amputated worms at 4 dpa. Numbers at left corner indicate the fraction of parasites exhibiting similar expression patterns, as determined from at least two biological replicates. Scale bar: 100 µm.

Previous studies indicate that a majority of neoblasts in adult schistosomes are destined to give rise to new tegumental cells (*8*). As puncture, amputation, or PZQ-mediate injuries are likely to damage not only the tegument but other worm tissues, we reasoned there may be molecular changes in the neoblast pool as these stem cells shift their fates away from the tegument and toward other tissue types. One prominent example is the gene *eled*, which is expressed in a large number of juvenile neoblasts (*11*) but becomes restricted to a small population of gut-associated neoblasts in adults (*9*). Consistent with the model that injury results in molecular remodeling of the neoblasts, both poke and amputation injuries induced marked increases in the number of *eled*^+^ cells around sites of injury, and sublethal PZQ exposure led to a significant increase in *eled*^+^ cells throughout the entire trunk (Fig. 1A bottom). These injury-responsive *eled*^+^ cells are predominantly EdU^+^, indicating these cells are proliferative (Fig. S1A). Moreover, *eled*^+^ and EdU^+^ cell doublets were observed following injury (Fig. S1A), indicating cell division occurred (*7*).

To examine how neoblasts respond to injury over time, we monitored neoblast proliferation in worms over a period of 1-5 days post-injury. We observed proliferative responses starting at day 2 post-injury, peaking between day 3 and 4, followed by a decline at day 5 (Fig. S1B-E). Intriguingly, we observed an interesting pattern of *eled*^+^ cell expression following amputation. At day 2 post-amputation injury, the *eled*^+^ response was localized to the injury site in both the anterior and posterior trunks (Fig. 1B). By day 3, *eled*^+^ cells had expanded in the amputated anterior trunk, while the response in the amputated posterior trunk remained localized (Fig. 1B). Notably, this anterior-driven response was independent of the exact amputated location. Whether the amputation occurred at the very anterior trunk or at the tail, the response in the anterior trunk remained consistent (Fig. 1C). Additionally, the presence of the head was not required for this expanded *eled* response, as trunk fragments lacking both head and tail still maintained elevated *eled* expression, whereas this was not observed in worms subjected to head amputation alone (Fig. 1D). Altogether, our results showing both mechanical and chemical injury induces proliferative responses in adult worms, but with different spatial patterns depending on the nature and location of the wound.

### A *bHLH* transcription factor responds to mechanical injury in adult *S. mansoni*

To identify factors regulating the proliferative response following injury, we performed bulk RNA-seq on posterior trunk fragments collected at 4 days post-amputation (Fig. 2A). Male worms were subjected to either tail-only or tail-and-head amputation, and posterior fragments were selected because they exhibit robust proliferation while minimizing contribution from the testes that express high levels of *eled* and other germline associated genes (Fig. 1D). We detected 507 and 483 differential expressed genes (DEGs) (log2FC > 0.5, *P*adj < 0.05) in tail-amputated worms and tail-and-head-amputated worms, respectively (Data S1 and S2). Both amputated groups shared highly significant up-regulated DEGs (log2FC > 1.5) (Fig. S2A, Data S3), suggesting the head is not a major contributor to these injury-induced responses, consistent with the observed proliferative responses (Fig. 1D).

**Figure 2.**
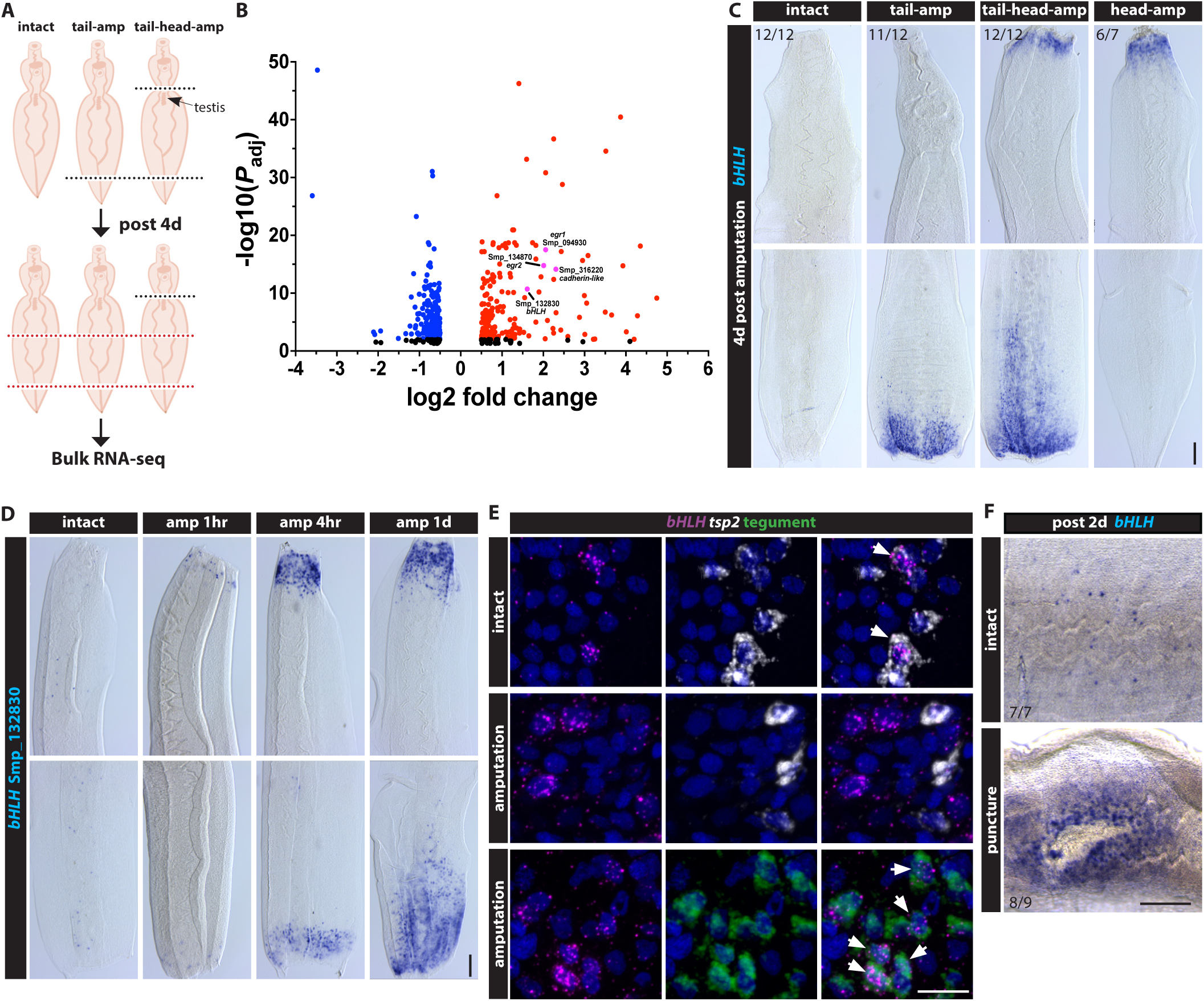
***bHLH* is mechanical injury responsive.** (A) Schematic of the bulk RNA-seq strategy on amputated worms. Male parasites were amputated either the tail only or both the tail and head. At 4 days post-amputation, the posterior trunk fragments (highlighted with dashed red lines) were collected for bulk RNA-seq analysis. (B) Volcano plot showing 483 differentially expressed genes (DEGs) (Log2FC>0.5, *P*adj<0.05) in tail-and-head-amputated worms compared to intact controls. Blue and red dots indicate significantly down-regulated and up-regulated DEGs (*P*adj<0.01), respectively. Pink dots highlight genes of interest. (**C**) WISH showing localized *bHLH* (Smp_132830) expression in tail-, tail-and-head- and head-amputated worms at 4 days post-amputation (dpa). Numbers in upper-left corner indicate the fraction of parasites exhibiting similar expression patterns, as determined from at least one biological replicate. Scale bar: 100 µm. (**D**) WISH time-course showing rapid induction of *bHLH* at 1 hour, 4 hours, and 1 day post amputation in tail-and-head-amputated worms. Representative images are shown for 1hr (intact: n=12/12 males, amputation: n =8/10 males), 4hr (intact: n=12/12, amputation: n=13/13), 1d (intact: n=9/9, amputation: n=8/8). Data were collected from at least three biological replicates. Scale bar: 100 µm. (**E**) Double FISH in intact and 4 dpa worms showing *bHLH* expression relative to the tegument progenitor marker *tsp2* and tegument marker *calpain*. In intact worms, *bHLH* is primarily detected in *tsp2*^+^ cells, whereas amputation-responsive *bHLH* shows marked upregulation and shifts predominantly to the tegument cells. n>5 parasites per group from two biological replicates. Scale bar: 50 µm. (**F**) WISH showing *bHLH* induction at 2 days post-puncture injury. Numbers in the bottom-left corner indicate the fraction of parasites exhibiting similar expression patterns, as determined from two biological replicates. Scale bar: 100 µm.

Among these DEGs, we identified genes encoding several transcription factors and transmembrane proteins (Fig. 2B), including a *bHLH* (basic helix loop helix domain containing protein, Smp_132830), a pair of *egr*-like proteins (early growth response protein, Smp_094930 and Smp_134870), and a cadherin-like protein (cadherin-like domain containing protein, Smp_316220), all of which responded to amputation (Fig. 2C, Fig. S2B). Notably, the *bHLH* and *egr2* exhibited a rapid response to injury, showing weak responses 1hr after injury and robust localized responses 4hr after injury at the amputated sites (Fig. 2D, Fig. S2C). This contrasts with the expression of the *cadherin-like* gene, whose induction was only observed 1d after injury (Fig. S2D). Above all, these injury-induced factors were activated substantially earlier than changes in cell proliferation, suggesting their potential role in regulating the proliferative response.

In planarians, the regenerative microenvironment is highly dynamic and undergoes spatial and transcriptional reorganization in response to injury (*12, 13*). Given that injury also induces molecular remodeling of neoblasts in adult worms (Fig. 1A), we next asked whether these injury-responsive factors exhibit similar cell-type specific changes in expression. To define the cellular identity of these genes, we examined their expression relative to tissue-specific markers. In intact worms, *bHLH* was primarily expressed in the tegument progenitor *tsp2*^+^ cells (Fig. 2E), consistent with its abundance in scRNA-seq data from intact adult worms (Fig. S3A) (*14*). However, amputation-responsive *bHLH* exhibited a marked increase in expression and was predominantly localized to tegument cells (Fig. 2E). This robust injury-induced expression was absent from *tsp2*^+^ cells (Fig. 2E), neoblast progeny (Fig. S3C) or proliferative cells (Fig. S3D, E), despite the prediction of intact adult scRNA-seq data (Fig. S3A). A similar amputation-induced shift was observed for *egr2*, which localized to tegument cells of intact worms (Fig. S4A, B) but expressed primarily in muscle cells following injury (Fig. S4C). Likewise, *cadherin-like* was rarely detected in intact worms, but amputation-responsive *cadherin-like* was primarily localized to tegument cells (Fig. S4D, E). Altogether, amputation-responsive factors exhibited marked shifts in cellular localization following injury. Additionally, we also observed that while *bHLH* responded similarly to puncture injury (Fig. 2F, Fig. S5A) but not to PZQ-triggered injury (Fig. S5B, C), suggesting that *bHLH* expression may be specifically tied to the mechanical injury response and that chemical-induced injury is interpreted by alternate mechanisms.

### Injury-responsive *bHLH* is required for cell proliferation and new parenchyma production following mechanical injury

To explore the role of injury-responsive factors in regenerative responses, we performed *in vitro* RNAi for 14 days prior to amputation and quantified injury-responsive proliferation at 4 days post amputation (Fig. 3A). Consistent with previous observations, amputation significantly increased the number of EdU^+^ and *eled*^+^ cells in control RNAi parasites (Fig. 3B top and Fig. 3C, D). In contrast, *bHLH* RNAi worms showed no significant difference between intact and amputated worms (Fig. 3B bottom, Fig. 3C, D) and exhibited significantly fewer EdU^+^ and *eled*^+^ cells compared to amputated controls (Fig. 3C, D). A similar reduction in injury-responsive proliferation was also observed following puncture injury (Fig. 3E-G). Together, these results demonstrate *bHLH* is required for mechanical injury-induced cell proliferation.

**Figure 3.**
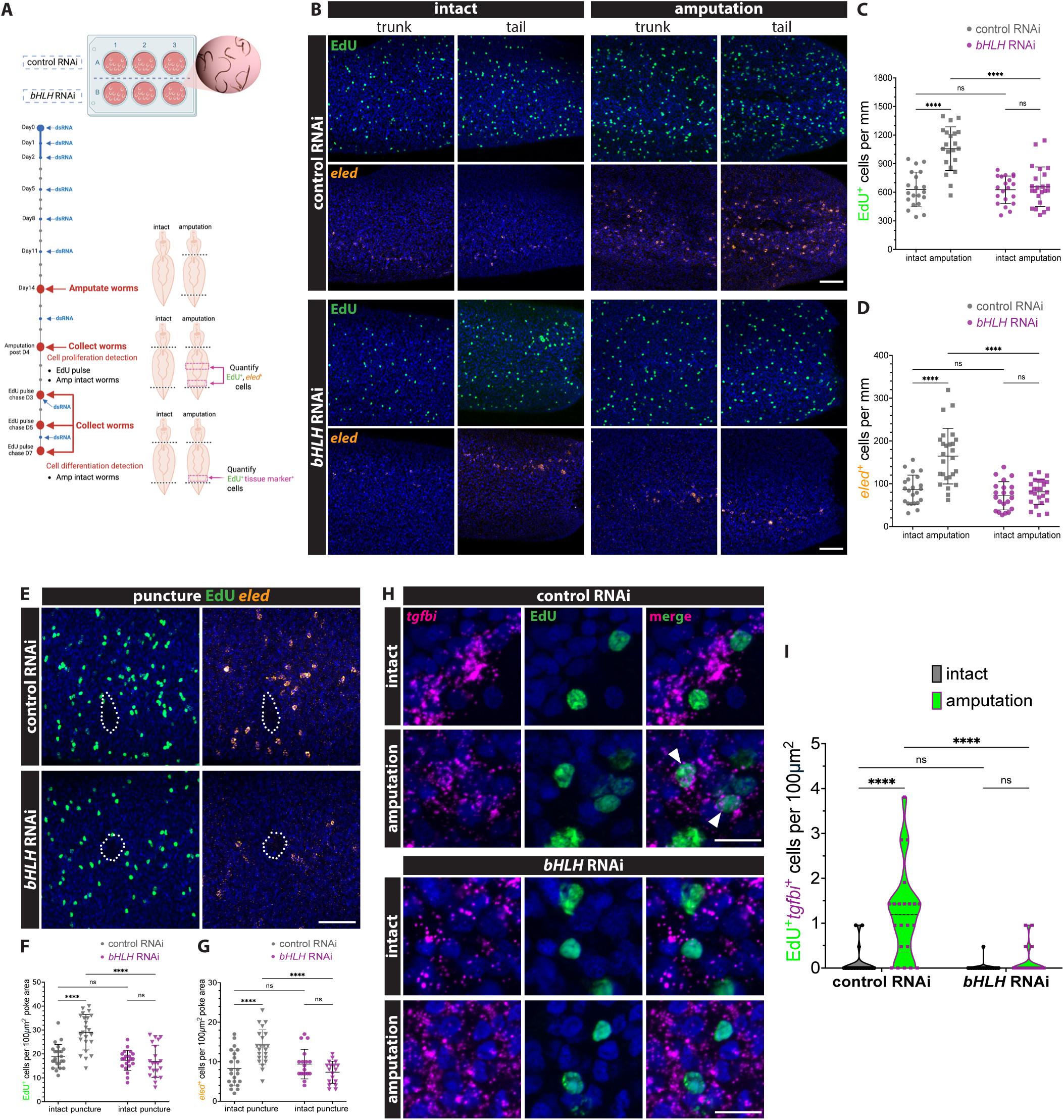
Injury-responsive *bHLH* is required for cell proliferation and new parenchyma production following mechanical injury. **(A)** Schematic illustrating the RNAi strategy used to assess *bHLH* function in injury-induced cell proliferation and differentiation. Male worms were subjected to *in vitro* RNAi for 14 days prior to amputation, with RNAi treatment continuing for an additional 4 days post-amputation (dpa). At 4 dpa, worms were pulsed with EdU and collected for proliferation analysis, or harvested after 3-, 5- and 7-days chase period for differentiation analysis. (**B**) FISH showing EdU^+^ and *eled*^+^ cells in trunk regions and at amputated sites of intact and amputated worms at 4 dpa following control RNAi or *bHLH* RNAi. (**C-D**) Quantification of (C) EdU^+^ and (D) *eled*^+^ cells. n>20 parasites per group from three biological replicates. Each data point represents the average number of EdU^+^ or *eled*^+^ per worm, calculated from combined trunk and tail sections. (**E**) FISH showing EdU^+^ and *eled*^+^ cells at 3 days after puncture injury following control RNAi or *bHLH* RNAi. (**F-G**) Quantification of (F) EdU^+^ and (G) *eled*^+^ cells. n>18 parasites per group from three biological replicates. Each data point represents the number of EdU^+^ or *eled*^+^ cells per 100 µm^2^ area surrounding the puncture site. (**H**) FISH for parenchymal marker (*tgfbi*) with EdU detection at Day 7 following an EdU pulse in control RNAi and *bHLH* RNAi worms. Arrows indicate EdU^+^ parenchymal cells. (**I**) Violin plot with individual dots showing quantification of EdU^+^*tgfbi*^+^ cells per 100 µm^2^ at amputated sites following a 7-day chase. n>20 parasites per group from three biological replicates. Scale bar: 50 µm. *** indicates *P*<0.001, ns indicates non-significant.

Because injury increased neoblast numbers, we next considered two possible fates for these cells: either they remain restricted to tegument and gut lineages, or they acquire a broader developmental potential capable of generating additional cell types. To distinguish between these possibilities, we performed *in vitro* EdU pulse-chase assays to track neoblast differentiation into multiple cell types following amputation, including tegument progenitor cells (*tsp2*^+^), mature tegument cells, gut cells, parenchymal cells, muscle cells and neuronal cells (Fig. 3A). In intact worms, a small number of new tegument progenitor (*tsp2*^+^) cells were detected, with a gradual increase from day 3 to 5 post-pulse (Fig. S6). Amputation significantly increased the production of *tsp2*^+^ cells (Fig. S6) and mature tegument cells (Fig. S7), particularly at early time points (Fig. S7A-D). Amputation also increased the generation of new gut cells (Fig. S8A-C), indicating an overall increase in tegument and gut lineage output following injury; However, these effects were not significantly altered by *bHLH* knockdown (Fig. S7, 8A-C). In contrast, newly generated parenchymal cells were rarely detected in intact worms, whereas amputation robustly induced their production after a 7-day chase (Fig. 3H top, 3I). This increase was largely abolished in *bHLH* RNAi worms, which produced very few new parenchymal cells following amputation (Fig. 3H bottom, 3I). Notably, amputation did not significantly induce the production of new muscle or neuronal cells under either condition (Fig. S8D-F). Collectively, our results demonstrate that *bHLH* is required for injury-induced cell proliferation and specifically for the generation of new parenchymal cells during regeneration.

### *wd40* acts downstream of *bHLH* to regulate injury-induced cell proliferation

To further identify downstream effectors of *bHLH* that mediate injury-induced proliferation, we performed bulk RNA-seq on amputated worms following *bHLH* RNAi (Fig. 4A). Transcriptomic comparison between amputated *bHLH* RNAi and control RNAi worms identified genes potentially acting downstream of *bHLH* (Fig. 4B, Data S4). A subset of these differentially expressed genes (DEGs) was selected for functional RNAi screening to assess their roles in injury-induced proliferation. This screen identified two wd40 domain-containing proteins (*wd40*), Smp_125760 (*wd40-1*) and Smp_054610 (*wd40-2*), as amputation-responsive genes (Fig. 4C, Fig. S9A, B). Knockdown of *wd40-2* impaired the injury-induced increase in EdU^+^ and *eled*^+^ cells (Fig. 4D-F), recapitulating the proliferative defect observed upon *bHLH* knockdown (Fig. 3B-D), although the two genes were not spatially co-localized (Fig. S9C).

**Figure 4.**
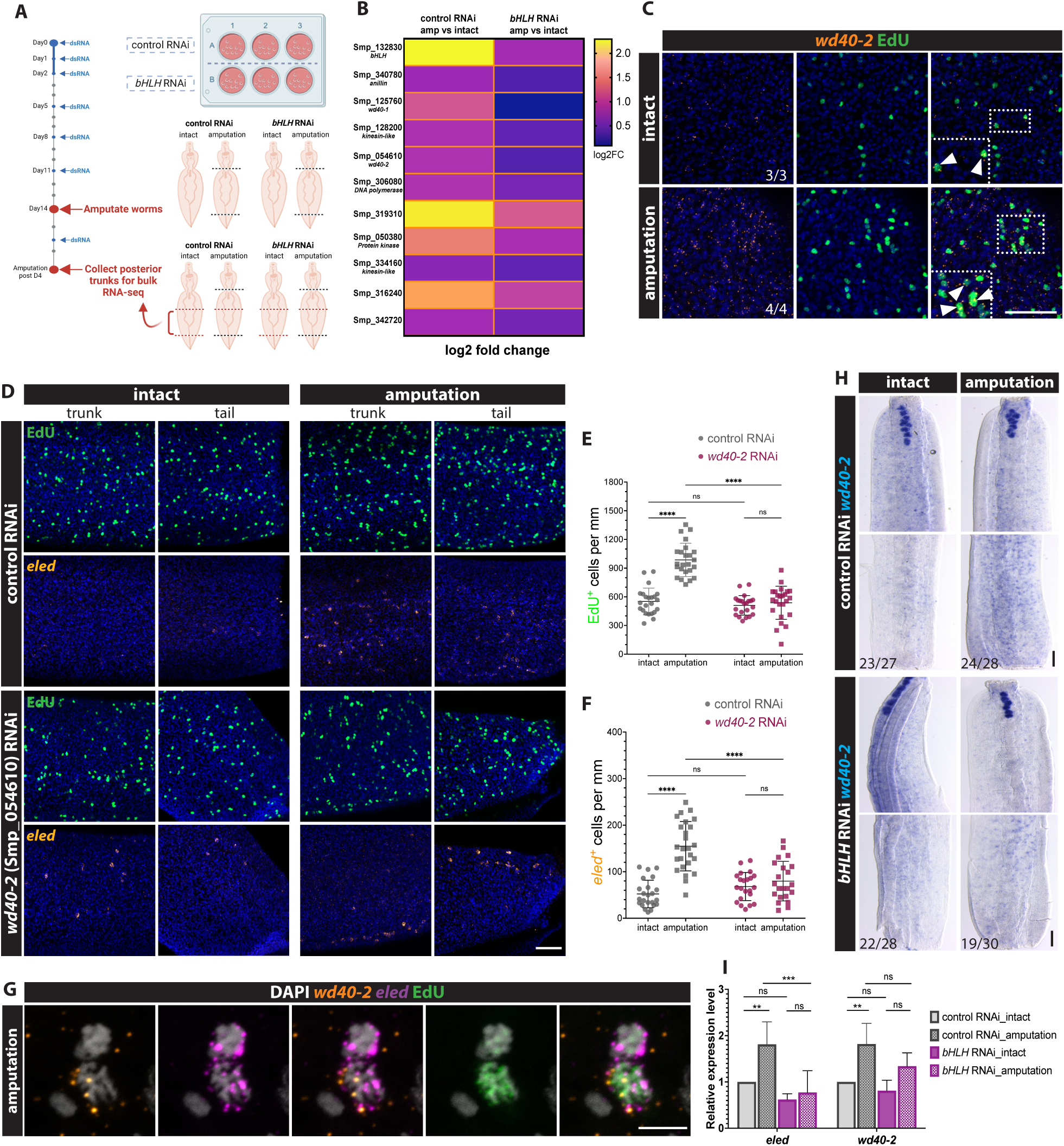
Amputation-responsive *wd40-2* acts downstream of *bHLH* to regulate injury-induced cell proliferation. **(A)** Schematic illustrating the *bHLH* RNAi strategy for bulk RNA-seq following amputation. Male worms were subjected to *in vitro bHLH* RNAi for 14 days prior to tail-and-head amputation, with RNAi treatment continuing for an additional 4 days post-amputation (dpa). At 4 dpa, the posterior trunk fragments from four conditions (control or *bHLH* RNAi; intact or amputated) were collected for bulk RNA-seq analysis. (**B**) Heatmap of 11 candidate genes significantly up-regulated following amputation in control RNAi worms (*P*adj<0.001), but not in *bHLH* RNAi worms (*P*adj>0.05). (**C**) FISH showing *wd40-2* expression with EdU detection at 4 dpa. The dotted box is shown in higher magnification (bottom-left); arrows indicate *wd40-2*^+^EdU^+^ cells. Numbers at corner indicate the fraction of parasites with similar expression patterns. Scale bar: 50 µm. (**D**) FISH showing EdU^+^ and *eled*^+^ cells in trunk regions and at amputated sites of intact and amputated worms at 4 dpa following control RNAi or *wd40-2* RNAi. Scale bar: 50 µm. (**E-F**) Quantification of (E) EdU^+^ and (F) *eled*^+^ cells. Each data point represents the average number of EdU^+^ or *eled*^+^ per worm, calculated from combined trunk and tail sections. n>22 parasites per group from three biological replicates. (**G**) Double FISH showing *wd40-2* was co-localized with *eled* within the same EdU*^+^*cell in amputated worms at 4 dpa. n>6 parasite from two biological replicates. Scale bar: 5 µm. (**H**) WISH showing *wd40-2* expression in both intact and amputated worms following *bHLH* RNAi. Numbers at left corner indicate the fraction of parasites with similar expression patterns, from 4 biological replicates. Scale bar: 100 µm. (**I**) qPCR quantification of *eled* and *wd40-2* expression in intact and amputated worms following *bHLH* RNAi, data are representative of three biological replicates (n>8 parasites per group). ** indicates *P*<0.01, *** indicates *P*<0.001, ns indicates non-significant.

Notably, *wd40-2* knockdown did not affect steady-state neoblast numbers but attenuated the injury-induced proliferative increase (Fig. 4D-F). In contrast, *wd40-1* knockdown reduced overall proliferative cell numbers, indicating a requirement for maintaining the proliferative population (Fig. S9D-F). Consistent with these results, both *wd40* genes were predominantly expressed in proliferative cells (Fig. 4C, Fig. S9B). In addition, *wd40-2* was co-expressed with amputation-responsive *eled* in the same cells (Fig. 4G), indicating *wd40-2* functions within the *eled*^+^ cell population. Importantly, suppression of injury-responsive *bHLH* reduced amputation-induced expression of both *wd40* genes (Fig. 4H, I and Fig. S9G) placing *wd40* genes downstream of *bHLH* in the injury-induced proliferative program. Together with the phenotypic analyses, these findings identify *wd40* genes as key effectors linking *bHLH* activity to regenerative cell proliferation.

## Discussion

Adult schistosomes persist within the hostile environment of the host vasculature for decades, yet their regenerative response to tissue damage remains poorly understood. Here we show that adult *Schistosoma mansoni* trigger robust proliferative responses to multiple forms of injury. Mechanical injury rapidly induces the transcription factor *bHLH*, which undergo striking shifts in cellular localization following amputation. The expression of *bHLH* becomes enriched in tegumental cells, suggesting that differentiated tissue may establish a transient regenerative microenvironment that promotes injury-induced proliferation of *eled* neoblasts (Fig. 5, top). Functional analyses further reveal that *bHLH* is required for injury-induced neoblast proliferation and the production of new parenchymal cells (Fig. 5, bottom), identifying a mechanical injury– responsive transcriptional program that links tissue damage to neoblast activation.

**Figure 5.**
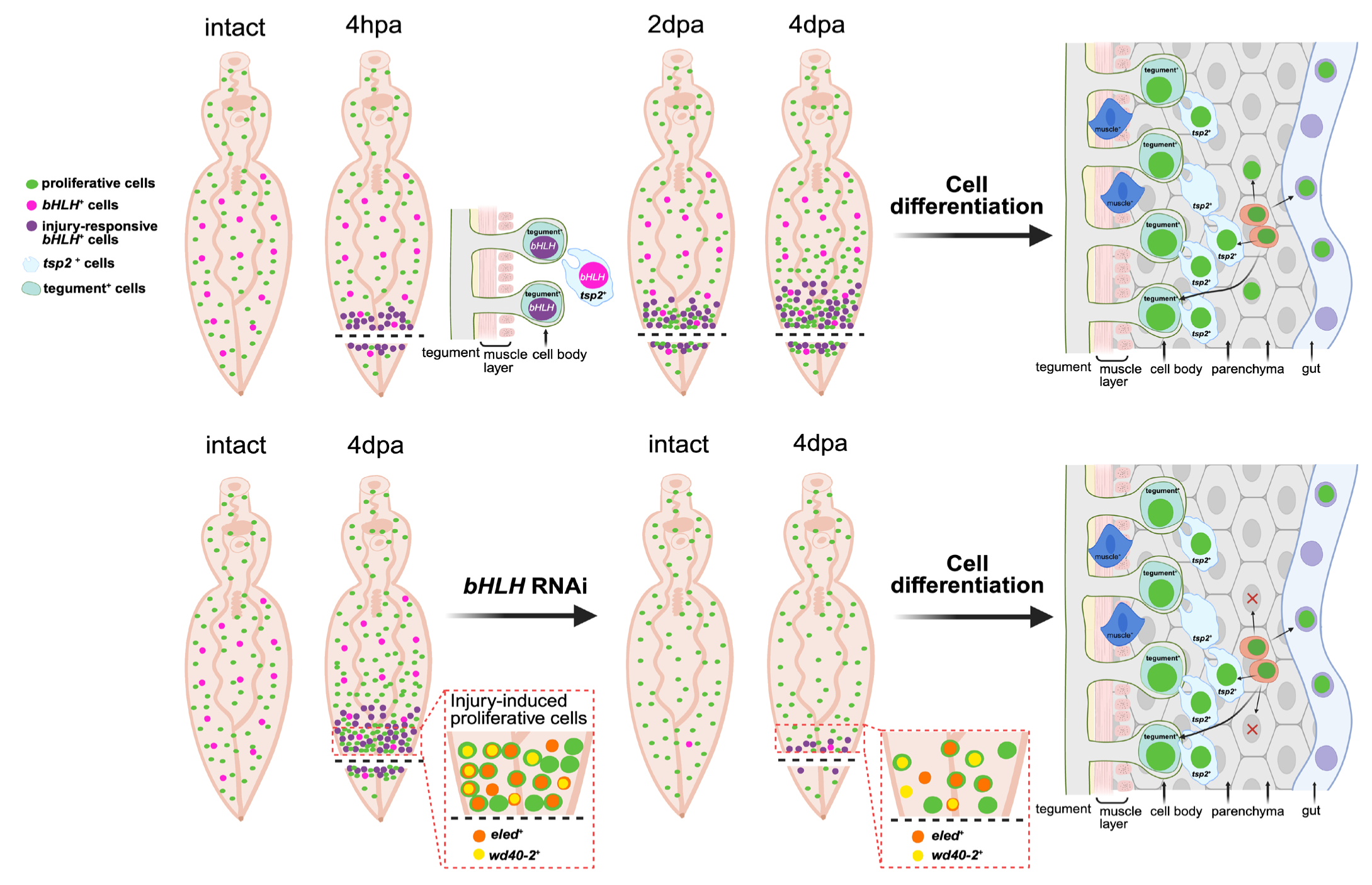
Regenerative model of a mechanical injury-activated microenvironment–stem cell program in adult *S. mansoni*. **(A)** (**Top**) Within 4 hours post-amputation (4 hpa), transcription factor *bHLH* is rapidly induced and re-localizes, shifting from *tsp2*^+^ cells to tegument cells. The differentiated tissue contributes to a transient regenerative microenvironment that drives a localized proliferative response at 2 days post-amputation (2 dpa), which expands anteriorly within the trunk by 4 days post-amputation (4 dpa). Injury-induced neoblasts give rise to tegument progenitor *tsp2*^+^ cells, tegument cells, parenchymal cells and gut cells, but rarely generate muscle cells and do not produce neuronal cells. (**Bottom**) At 4 dpa, the amputation-responsive *eled*^+^ population co-expresses *wd40-2* gene in proliferative cells. Following *bHLH* knockdown, the regenerative microenvironment is disrupted, resulting in impaired injury-induced proliferation and reduced parenchymal cell production, while the generation of *tsp2*^+^, tegument, gut, muscle and neuronal cells is largely unaffected.

### Different types of injuries trigger distinct proliferative dynamics

Mechanical injuries (puncture and amputation) both initially induced localized proliferative responses near the wound site at 2 days post-injury (Fig. 1A). However, their later dynamics diverged. While the proliferation following puncture injury remained largely restricted to the injury site, amputation subsequently triggered a broader proliferative expansion at later time points (3-4 days post-injury) (Fig. 1B-D). This pattern suggests that schistosomes may prioritize rapid wound repair before engaging a broader systematic proliferative response. The proliferative pattern differs from that observed in regenerative flatworms such as planarian. After amputation, planarians exhibit a biphasic mitotic response consisting of an early systemic mitotic burst triggered by injury stress and a later localized proliferative peak associated with regeneration of missing tissues (*15, 16*). In contrast, schistosomes displayed only a single proliferative peak, and localized proliferation preceded the later widespread response (Fig. S1B–D). These differences likely reflect fundamental distinctions in regenerative capacity between parasitic and free-living flatworms. Meanwhile, chemical injury triggered a markedly different response. Sublethal exposure to PZQ induced a widespread proliferative response throughout the worm (Fig. 1A, Fig. S1D), likely reflecting systemic tissue stress. In contrast to mechanical injury, chemical damage did not induce *bHLH* expression (Fig. S5B, C), suggesting that distinct molecular pathways may govern responses to localized tissue damage versus systemic stress. Alternatively, because injury-responsive *bHLH* is expressed to the tegument (Fig. 2E), PZQ-induced disruption of tegument integrity may impair its activation (*6, 17*). Together, these results suggest that adult schistosomes deploy flexible injury-response programs that tailor stem cell activity to the nature of the injury.

### *eled* neoblasts represent a plastic injury-responsive population

Juvenile neoblasts exhibit expanded molecular diversity (*11*), suggesting they have broader developmental potential. Notably, *eled* neoblasts are widely distributed in juvenile worms and associated with rapid growth (*11*), whereas in adult worms they are sparse and largely restricted to the gut. Strikingly, we observed a marked expansion of *eled* cells following injury (Fig. 1A), with amputation inducing a pronounced anterior-biased accumulation (Fig. 1B), a spatial pattern that resembles the growth dynamics in juvenile parasites. These findings suggest injury may enable adult worms to transiently re-enter a juvenile-like proliferative state, thereby promoting tissue repair. Rather than representing a fixed lineage, the *eled* population function as a plastic stem cell pool that expands in response to increased regenerative demands, such as the increased production of parenchymal cells observed following amputation (Fig. 3H). In this context, injury-induced activation of developmentally associated stem cell programs may represent an adaptive strategy that supports the long-term persistence of these parasites within their hosts.

### Injury-induced transcriptional shifts establish a regenerative-microenvironment

Our results further suggest that injury induces a transient transcriptional “microenvironment” that regulates neoblast activation. Two transcription factors, *bHLH* and *egr2*, responded rapidly to mechanical injury and exhibited striking shifts in cellular localization following amputation. In intact worms, *bHLH* expression was primarily associated with *tsp2* cells; however, after injury its expression shifted predominantly to tegument cells (Fig. 2E). Similarly, *egr2* was detected in muscle cells following injury but was not expressed in these cells under homeostatic conditions (Fig. S4B, C). These injury-dependent changes in cellular identity suggest that differentiated tissues dynamically alter their transcriptional programs in response to damage.

Notably, neither *bHLH* nor *egr2* was expressed in proliferative neoblasts or neoblast progeny (Fig. S3 and S4A-C) indicating that these factors do not act directly within stem cells. Instead, our functional data suggest that *bHLH* regulates injury-induced proliferation indirectly through downstream targets, including wd40 repeat–containing protein 2 (*wd40-2*), which is expressed in *eled*^+^ proliferative cells. WD40-2 is homologous to WD repeat and high mobility group (HMG)-box DNA-binding protein 1 (WDHD1) (Fig. S10), a protein that plays critical roles in DNA replication, DNA repair and sister chromatid cohesion (*18–22*). These functions suggest that WD40-2 may support injury-induced proliferative response by promoting DNA replication and maintaining genome integrity in dividing cells. Together, these observations support a model in which injury-responsive transcriptional changes in differentiated tissues generate signals that promote neoblast proliferation, effectively establishing a local regenerative microenvironment (*12, 13*).

### Limited regenerative capacity of adult *S. mansoni*

Despite this regenerative response, adult schistosomes appear to possess limited lineage potential. Following injury, adult neoblasts produced increased numbers of tegument progenitors, as well as tegument cells and gut cells (Fig. S6-8), which was expected since intact worms continuously maintain these tissues and *bHLH* knockdown did not significantly impair their production (Fig. S6-8). Surprisingly, injured adult worms generated new parenchymal cells, a phenomenon not previously reported, and this response required *bHLH* (Fig. 3H). In contrast, we observed little production of new muscle or neuronal cells (Fig. S8D-F). This selective production raises the possibility that adult schistosome regeneration is lineage-restricted, with neoblasts preferentially generating specific cell types during tissue repair. Alternatively, other neoblast populations may exist that restore a more juvenile-like proliferative state capable of repairing muscle and neuronal tissues. Such targeted repair may be sufficient to maintain tissue integrity under host-imposed stresses without engaging large-scale regeneration.

## Conclusions

Targeting components of this microenvironment−neoblast axis, such as *bHLH* or its downstream effectors, could represent a strategy to compromise parasite resilience and enhance anti-schistosome therapies. In our study, injury-induced *wd40* expression was only partially suppressed following *bHLH* knockdown, likely reflecting incomplete RNAi efficiency, as amputated worms retained low levels of *bHLH* expression (Fig. S11). These findings highlight the importance of identifying upstream regulators that activate *bHLH* following mechanical injury. Furthermore, because WD40 proteins often function as scaffolds that assemble signaling complexes (*23*), identifying factors that cooperate with WD40 may reveal how microenvironment-derived injury signals are integrated with intrinsic neoblast proliferation programs. Lastly, determining whether this injury-responsive axis contributes to parasite persistence within the host will be an important next step.

Together, these findings suggest that adult schistosomes respond to tissue damage by transiently reprogramming differentiated tissues into a regenerative microenvironment that activates neoblast proliferation, revealing a previously unrecognized strategy that may contribute to the extraordinary longevity of these parasites within their hosts.

## Materials and methods

### Parasite acquisition and culture

Adult *S. mansoni* (NMRI strain, 6–7 weeks post-infection) were obtained from infected female mice by hepatic portal vein perfusion with 37°C DMEM (Sigma-Aldrich, St. Louis, MO) plus 10% Serum (either Fetal Calf Serum or Horse Serum) and heparin. Parasites were cultured as previously described (*8*). All experiments were performed with male parasites to maximize the amount of somatic tissue present. Experiments with and care of vertebrate animals were performed in accordance with protocols approved by the Institutional Animal Care and Use Committee (IACUC) of UT Southwestern Medical Center (approval APN: 2017-102092).

### Worm Injury

For mechanical injury, worms were anesthetized in a 0.25% solution of the ethyl 3-aminobenzoate methanesulfonate (Sigma-Aldrich, St. Louis, MO) dissolved in Basch Media 169. For puncture injuries, worms were gently pipetted onto the surface of a 35 mm Petri dish containing solidified 4% agarose prepared in H_2_O. After excess liquid was removed, individual worms were perforated with a sharpened tungsten needle. The impaled parasites were then carefully transferred into fresh media using a pipette tip. As a control, “mock-injured” parasites were similarly transferred to Petri dishes but were not subjected to needle injury. For amputation injury, half of the worms in each well were amputated at define anatomical locations using a sharpened probe in 6 well-plates, while the remaining worms were left uncut as “intact controls”. After the procedure, the anesthetic medium was replaced with fresh Basch Media 169. For PZQ-induced injury, worms were treated with 1μg/mL PZQ (Selleckchem, Houston, TX) dissolved in ethanol for 6 hours, while control worms received an equivalent volume of ethanol. Following treatment, the medium was replaced with fresh Basch Media 169.

### RNA interference

20 freshly perfused male parasites were placed into 6-well plates and cultured in 10mL in Basch Media 169 supplemented with 30 μg/mL dsRNA for 14 days. dsRNA was generated by *in vitro* transcription and replaced with fresh media on Day 0, 1, 2, 5, 8, 11. On day 14, the worms were anesthetized to amputate both the head and tail, then refreshed with Basch Media 169 with dsRNA. The worms were cultured *in vitro* for 4 additional days, with one media/dsRNA change. At 4 days post-amputation, the worms were pulsed with 10 µM EdU for 4 hours before being fixed as previously described (*7*); for EdU pulse-chase experiments, the worms were pulsed with 10 µM EdU for 4 hr after which the media was changed, the worms were fixed following 3, 5 and 7-day chasing period. As a negative control for RNAi experiments, we used a non-specific dsRNA containing two bacterial genes (*24*). cDNAs used for RNAi and *in situ* hybridization analyses were cloned as previously described (*24*); oligonucleotide primer sequences are listed in Data S5.

### Parasite labeling and imaging

Colorimetric and fluorescence *in situ* hybridization detections were performed as previously described (*7, 8, 25*). EdU detection was performed as previously described (*7*). All fluorescently labeled parasites were counterstained with DAPI (1 μg/mL), cleared in 80% glycerol, and mounted on slides with Vectashield (Vector Laboratories).

Brightfield images were acquired on a Zeiss Axio Zoom V16 equipped with a transmitted light base and a Zeiss AxioCam 105 Color camera. Confocal imaging of fluorescently labeled samples was performed using either a Zeiss LSM900 or a Nikon A1 Laser Scanning Confocal Microscope. For cell quantification, cells were manually counted in maximum-intensity projections derived from confocal stacks. To determine the total number of labeled cells throughout the entire depth of the parasite (e.g. *eled^+^* and EdU^+^ cells), confocal stacks were collected and cell counts were normalized to the length of the imaged region in mm for amputation and PZQ-induced injury experiments; for poke injury, confocal stacks were acquired and a 100 µm^2^ area surrounding the injury site was selected for cell counting. To determine the percentage of EdU^+^ cells, images were acquired at amputation sites using an oil-immersion objective. For EdU^+^ parenchymal cell quantification, EdU^+^*tgfbi*^+^ cells were counted throughout the full depth of the worms and normalized to 100 µm^2^; for EdU^+^ tegument progenitor analysis, completed confocal stacks were collected, and both EdU^+^*tsp2*^+^ cells and total *tsp2*^+^ were counted. For EdU^+^ cells in the tegument, gut and muscle, counts were restricted to a 5 µm depth from the worm due to high signal intensity at amputation sites; For EdU^+^ gut cell quantification, the EdU^+^ intestinal nuclei^+^ and intestinal nuclei^+^ cells were counted.

### qPCR and RNA-seq analysis

Whole parasites were collected and homogenized in Trizol, and RNA extraction, cDNA preparation and qPCR were performed as previously described (*26*). Oligonucleotide primer sequences used for qPCR are listed in Data S5. For RNA-seq analysis, three biological replicates were performed for each condition. The samples were prepared by Illumina TruSeq stranded mRNA library kit. All samples were sequenced with one flow cell on Illumina NextSeq 550 sequencer with 75bp read lengths. Reads were mapped with STAR (v2.7.10a) (*27*) and *S. mansoni* genome sequence (v10) and GTF files used for mapping were acquired from Wormbase Parasite (*28*). Differential gene expression were performed with DESeq2 (version 1.46.0) (*29*). Raw and processed data have been deposited in NCBI (GSE339314 and GSE341215). The volcano plot in Fig. 2B was made with plotting log2 fold change expression (log2FC>0.5) and -log10 (*P*adj) of differential expressed genes (*P*adj < 0.05) and the heatmap in Fig. S2A was made with plotting log2 fold change of DEGs (log2FC>1.5, *P*adj<0.0001) in GraphPad Prism. For identifying injury-induced genes suppressed by *bHLH* RNAi, the DEGs (*P*adj<0.001) from control RNAi (amputated vs intact) and *bHLH* RNAi (amputated vs intact) comparisons were filtered and compared. DEGs that were significantly up regulated in control RNAi worms (*P*adj<0.001) but not in *bHLH* RNAi worms (*P*adj>0.05) were selected for further RNAi screening. The heatmap in Fig. 4B was made with plotting log2 fold change of candidate genes in GraphPad Prism. The Uniform manifold approximation plots (UMAP) plots were generated in R (version 4.4.2) using the adult scRNA-seq dataset (GSE146736_adult_scseq_seurat) (*9*). Gene expression was visualized using the Seurat FeaturePlot function with parameters min.cutoff =0 and order = TRUE.

### Statistical analysis

GraphPad Prism software (v11.0.2) processed and presented the data as the mean with SD. All two-way comparisons were analyzed using two-way analysis of variance (ANOVA), followed by pairwise comparison using Fisher’s Least Significant Difference (LSD) test. qPCR result in Fig. S5C was analyzed using paired t-test.

## Acknowledgments

The Schistosome Infected mice and *B. glabrata* snails were provided by the National Institute of Allergy and Infectious Diseases (NIAID) Schistosomiasis Resource Center of the Biomedical Research Institute (Rockville, MD, USA) through National Institutes of Health (NIH)-NIAID Contract HHSN272201700014I for distribution through BEI Resources. RNA-seq was performed with the aid of the Genomics Sequencing Core at the University of Texas Southwestern Medical Center (UTSW). Biorender was used for the schematic material in Figure 3A, 4A and Figure 5. We thank George Wendt for valuable comments and suggestions, and Jennifer Silverman for helpful feedback on this manuscript.

## Author contributions

Conceptualization and Supervision: J.J.C Methodology, Investigation, Visualization: L.Z Writing—original draft: L.Z Writing—reviewing & editing: L.Z and J.J.C

## Supplementary Materials

Figure S1 to S11 Data S1-S5

**Figure S1. Proliferative cell behavior following different types of injury in adult *S. mansoni*.**

(**A**) Double fluorescence *in situ* hybridization (dFISH) showing *eled*^+^ cells relative to EdU*^+^* cells. White dotted line indicates the approximate gut region. Cells within yellow dotted boxes are shown at higher magnification in the far-right panels. Yellow and red arrowheads indicate *eled*^+^ doublets that are EdU*^+^*/EdU^-^. Green arrowheads indicate *eled*^+^ cells within EdU*^+^* doublets, white arrowheads showed *eled*^+^ EdU^+^ cells. Scale bar: 10 µm. (**B-D**) FISH showing EdU*^+^* and *eled*^+^ cells at 1-5 days following (B) puncture, (C) amputation, and (D) PZQ-induced injury. Dotted line indicates the approximate sites of injury. Numbers represent the fraction of parasites exhibiting similar expression patterns, as determined from at least 1 biological replicate. Scale bar: 50 µm. (**E**) Quantification of EdU*^+^*(top) and *eled*^+^ (bottom) cells at 2-3 days post PZQ-induced injury. n>35 parasites per group from 4 biological replicates. * indicates *P*<0.05, ** indicates *P*<0.01, *** indicates *P*<0.001, **** indicates *P*<0.0001.

**Figure S2. Expression patterns of amputation-responsive candidate genes.**

(A) Heatmap illustrating significantly up-regulated genes (log2FC>1.5, *P*adj<0.0001) in tail-amputated and tail-and-head-amputated worms compared to intact controls. Red asterisk indicates the 4 genes of interest. (**B**) Left: Cartoon showing the worm’s anterior-posterior axis with head and tail amputation sites. Right: WISH showing localized expression of *egr1* (Smp_094930), *egr2* (Smp_134870) and *cadherin-like* (Smp_316220) at 4 days post amputation (dpa). (**C-D**) WISH time-course showing the induction of (C) *egr2* and (D) *cadherin-like* at 1 hour, 4 hours and 1 day post amputation. Numbers in the upper-left corner represent the fraction of parasites exhibiting similar expression patterns, as determined from at least 1 biological replicate. Scale bar: 100 µm.

**Figure S3. Amputation-responsive *bHLH* is not expressed in neoblasts or neoblast progeny.** (**A-B**) Uniform manifold approximation plots (UMAP) plots showing expression patterns of (A) *bHLH* and (B) Smp_194050 in adult schistosomes. Orange indicates high expression, blue indicates low/no expression. (**C-E**) Double FISH showing *bHLH* expression relative to the (C) neoblast progeny (Smp_194050^+^) cells, (D) *nanos2*^+^ cells and (E) *eled*^+^ cells, together with EdU labelling. Arrows indicate the co-localization of the signals. n>4 parasites per group from 1 biological replicate. Scale bar: 10 µm.

**Figure S4. *egr2* expression shifts from tegument to muscle following amputation.**

(**A**) Uniform manifold approximation plots (UMAP) plots showing expression patterns of *egr2* in adult schistosomes. (**B-C**) Double FISH showing *egr2* expression relative to (B) tegument (*calpain*^+^) cells and (C) muscle (*tpm2*^+^) cells in intact and amputated worms. In intact worms, *egr2* is expressed in tegument cells, whereas following amputation its expression shifts to muscle cells (intact: 2/117 *egr2*^+^muscle^+^ cells; amputation: 89/125 *egr2*^+^ muscle^+^ cells). (**D**) UMAP plots showing expression patterns of *cadherin-like* in adult schistosomes. (**E**) Double FISH showing *cadherin-like* expression relative to the tegument cells (top) and *bHLH*^+^ cells (bottom). n>4 parasite per group from 1 biological replicate. Scale bar: 50 µm.

**Figure S5. Mechanical injury-responsive *bHLH* does not responds to PZQ-induced injury.**

(**A**) FISH showing *bHLH* induction at 3 days post-puncture injury; data from 1 biological replicate. (**B**) FISH showing *bHLH* expression with EdU labelling at 3 days post PZQ-induced injury, numbers in the bottom-left corner represent the fraction of parasites exhibiting similar expression patterns from 2 biological replicates. (**C**) qPCR result showing *bHLH* expression at 3 days post PZQ-induced injury; data are representative of 3 biological replicates (n>5 parasites per group). Scale bar: 50 µm. * indicates *P*<0.05.

**Figure S6. Injury-responsive *bHLH* is not responsible for producing tegument progenitors following amputation.**

(**A**) FISH for tegument progenitor (*tsp2*^+^) cells with EdU detection at day 3 following an EdU pulse in control RNAi and *bHLH* RNAi worms. (**B**) Quantification of percentage EdU^+^*tsp2*^+^ cells after a 3-day chase. (**C-D**) (C) FISH and (D) quantification of tegument progenitor (*tsp2*^+^) cells at day 5 post-pulse in control RNAi and *bHLH* RNAi worms. Arrows indicate EdU^+^*tsp2*^+^ cells, percentages are shown in the upper-right corner. n>22 parasites per group from 3 biological replicates. Scale bar: 10 µm. ** indicates *P*<0.01, *** indicates *P*<0.001, **** indicates *P*<0.0001, ns indicates non-significant.

**Figure S7. Injury-responsive *bHLH* is not required for producing mature tegument following amputation.**

(**A**) FISH for tegument cells labelled with a tegumental marker cocktail (*calpain* Smp_214190, *npp-5* Smp_153390, *annexin* Smp_077720 and *gtp-4* Smp_105410) at day 3 post-pulse in control RNAi and *bHLH* RNAi worms. (**B**) Quantification of percentage EdU^+^tegumental cells after a 3-day chase. n>8 parasites per group from 2 biological replicates. (**C-D**) (C) FISH and (D) quantification of day 5 post-pulse. n>16 parasites per group from 3 biological replicates. (**E-F**) (E) FISH and (F) quantification of day 7 post-pulse. n>12 parasites per group from 2 biological replicates. Arrows indicate EdU^+^tegumental cells, percentages are shown in the upper-right corner. Scale bar: 10 µm. * indicates *P*<0.05, ** indicates *P*<0.01, *** indicates *P*<0.001, ns indicates non-significant.

**Figure S8. Injury-responsive *bHLH* is not required for producing gut following amputation.**

(**A**) FISH for gut (*cathepsin L*^+^) cells with EdU detection at 7 days post-pulse in control RNAi and *bHLH* RNAi worms. Arrows indicate EdU^+^ intestinal cells. (**B-C**) (B) Quantification of total EdU^+^intestinal cells and intestinal nuclei (n>18 parasite per group) and (C) percentage of EdU^+^intestinal cells from 3 biological replicates. * indicates *P*<0.05, ns indicates non-significant. (**D-E**) (D) FISH and (E) quantification for muscle (*tpm2*^+^) cells with EdU detection at 7 days post-pulse. Arrows indicate EdU^+^ muscle cells. Data are from 2 biological replicates (n>14 parasites per group). (**F**) FISH for neuronal (*7b2*^+^) cells with EdU detection at 7 days post-pulse. Numbers in the bottom-left corner represent the fraction of parasites exhibiting similar results within the replicate analyzed. Scale bar: 10 µm.

**Figure S9. *wd40-1* is required for maintenance of proliferative cells.**

**(A)** WISH showing *wd40-1* and *wd40-2* expression in worms at 4 days post amputation (dpa); data are from at least 2 biological replicates. Scale bar: 100 µm. (**B**) FISH showing *wd40-1* expression with EdU detection at 4 dpa. The dotted box is shown in higher magnification at bottom-left, arrows indicate *wd40-1*^+^EdU^+^ cells. Data are from one biological replicate. (**C**) Double FISH showing neither *wd40-1* nor *wd40-2* co-localizes with *bHLH* in amputated worms at 4 dpa (n>4 parasites from 1 biological replicate). (**D**) FISH showing EdU^+^ and *eled*^+^ cells in trunk regions and at amputated sites of intact and amputated worms at 4 dpa following *wd40-1* RNAi. (**E-F**) Quantification of (E) EdU^+^ and (F) *eled*^+^ cells. Each data point represents the average number of EdU^+^ or *eled*^+^ per worm, calculated from combined trunk and tail sections (n>23 parasites per group from 3 biological replicates). **** indicates *P*<0.0001. (**G**) FISH showing *wd40-1* expression in intact and amputated worms following *bHLH* RNAi (n>5 parasites from 1 biological replicate). Numbers at corner indicate the fraction of parasites exhibiting similar expression patterns. Scale bar (B-G): 50 µm.

**Figure S10. WD40-2 is homologous to WD40 repeat and high motility group-box DNA-binding protein1 (WDHD1).**

Protein sequence alignment of *Sm*WD40-2 with WDHD1 from *Schistosoma japonicum* (KAH8851080.1), *Clonorchis sinensis* (KAG5448950.1), *Homo sapiens* (XP_006720075.1), *Rattus norvegicus* (NP_001100725.1) and *Danio rerio* (XP_009291161.1). *Sm*WD40-2 is 1203 amino acids in length and contains WD40 repeat domains (highlighted in yellow rectangle), SepB domain (highlighted in green rectangle) and HMG box domain (highlighted in pink rectangle) (*22*).

**Figure S11. *bHLH* RNAi attenuates amputation-induced *bHLH* expression.**

WISH showing reduced but residual *bHLH* expression in worms at 4 days post amputation following *bHLH* knockdown. The signal observed in the gut of intact *bHLH* RNAi worms likely results from detection of ingested dsRNA, as worms were continuously incubated with dsRNA throughout the experiment. Numbers at corner indicate the fraction of parasites with similar expression patterns from one biological replicate. Scale bar: 100 µm.

**Data S1. Differentially expressed genes in adult *S. mansoni* following tail amputation.**

RNA-seq differential expression analysis comparing tail-amputated versus intact *S. mansoni* at 4 days post-amputation (dpa). The file contains three sheets: (1) all genes, with expression values and statistics from the differential expression analysis; (2) DEGs (*P*adj < 0.05); (3) DEGs (|log2FC|> 0.5, *Padj* < 0.05)

**Data S2. Differentially expressed genes in adult *S. mansoni* following both tail and head amputation.**

RNA-seq differential expression analysis comparing tail-and-head-amputated versus intact *S. mansoni* at 4 days post-amputation (dpa). The file contains three sheets: (1) all genes, with expression values and statistics from the differential expression analysis; (2) DEGs (*P*adj < 0.05); (3) DEGs (|log2FC|> 0.5, *Padj* < 0.05)

**Data S3. Most significant up-regulated Differentially expressed genes following tail-only or tail-and-head amputation in adult *S. mansoni*.**

The file contains three sheets: (1) DEGs (log2FC> 1.5, *Padj* < 0.00001) following tail**-only** amputation; (2) DEGs (log2FC> 1.5, *Padj* < 0.00001) following tail-and-head amputation; (3) **t**op 30 upregulated DEGs shared between tail-only and tail-and-head amputation.

**Data S4. RNA-seq differential expression analysis of *bHLH* RNAi versus control RNAi in *S. mansoni* following amputation.**

The file contains three sheets, sheet (1) and (2) listing all genes with expression values and statistics for the following comparisons: (1) control RNAi-amputation vs control RNAi-intact; (2) *bHLH* RNAi-amputation vs *bHLH* RNAi-intact; sheet (3) list of 11 candidate genes shown in Fig.4B: genes significantly up-regulated following amputation in control RNAi worms (log2FC>0, *P*adj<0.001), but not in *bHLH* RNAi worms (*P*adj>0.05). Expression statistics from both comparisons (control RNAi and *bHLH* RNAi) are shown side by side for each gene.

**Data S5. Oligonucleotides used in this study.**

The file contains two sheets: (1) Oligos used as templates for dsRNA synthesis or probe generation; (2) qPCR primers.

