## Supplementary figures and images for "An injury-responsive *bHLH* is required for regenerative responses following mechanical injury in adult *Schistosoma mansoni*"

### Figure S1

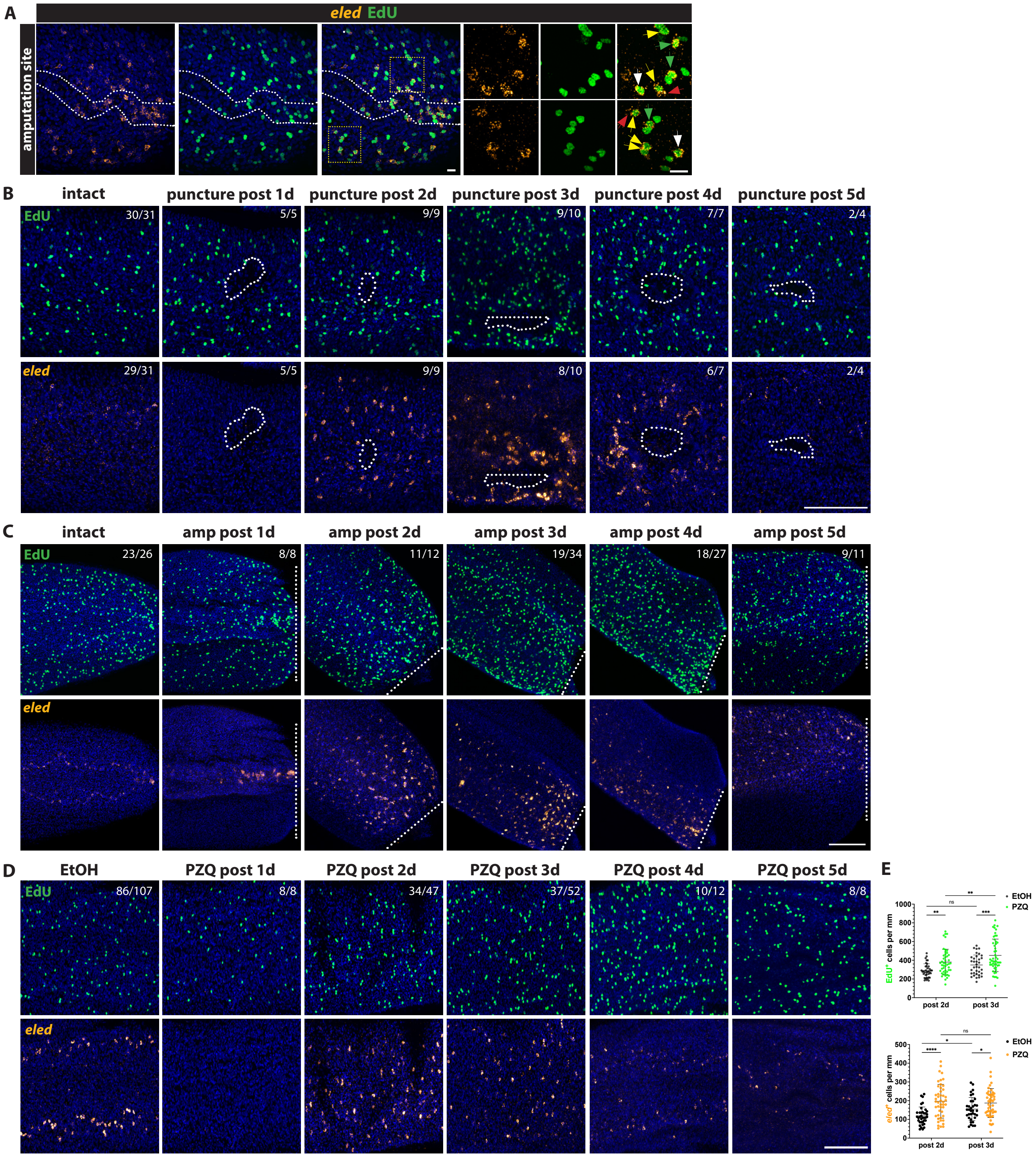

### Figure S2

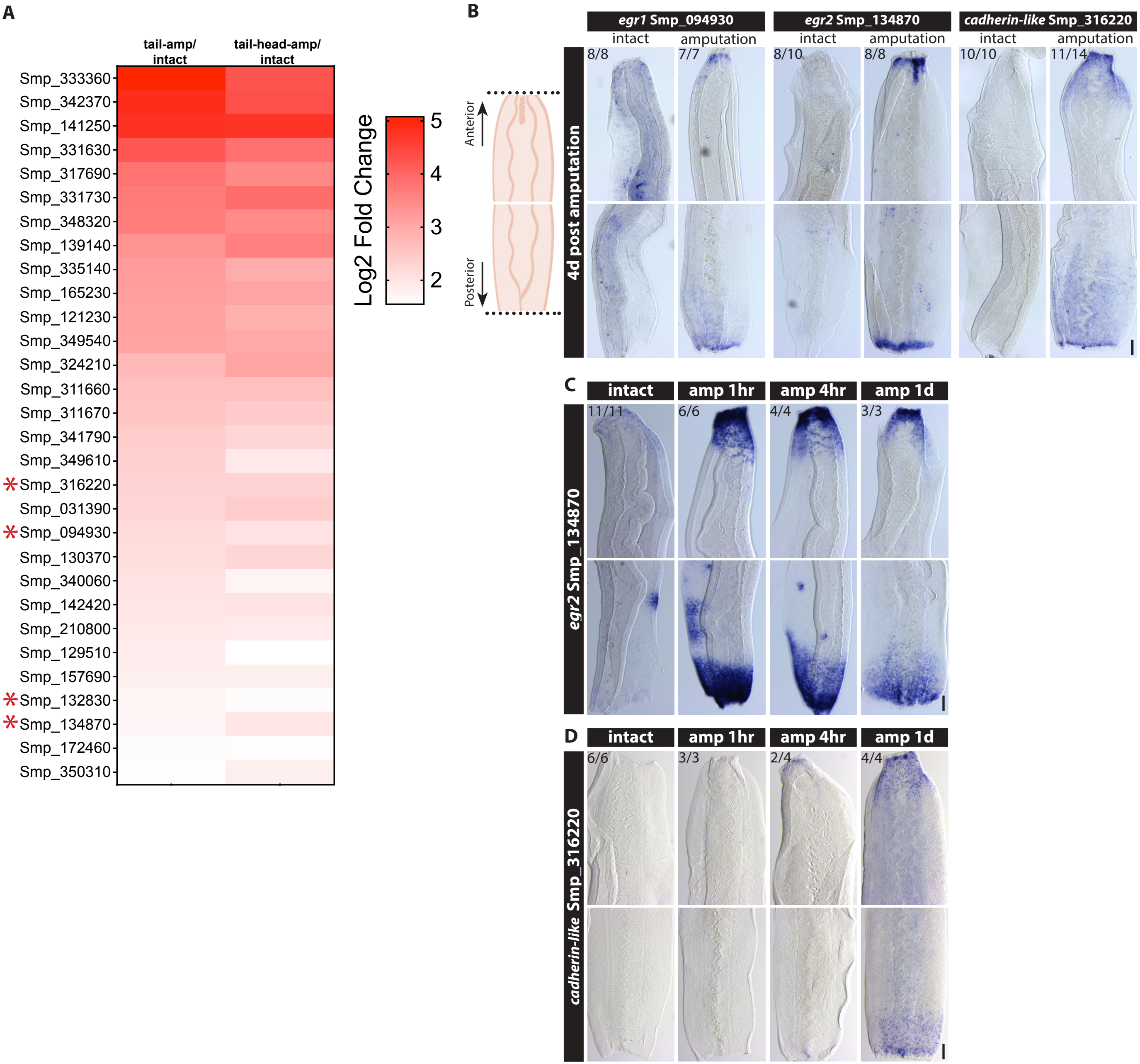

### Figure S3

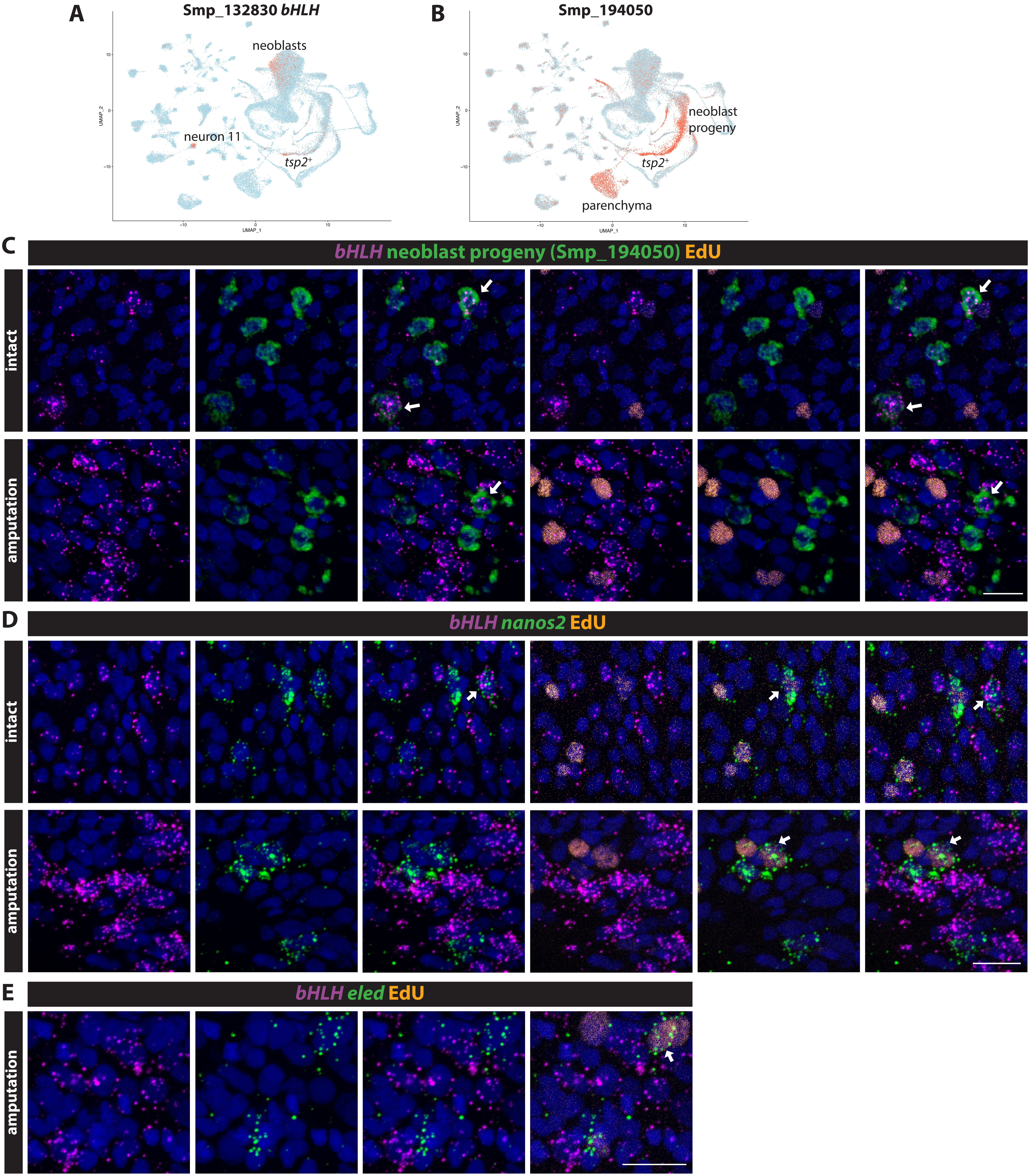

### Figure S4

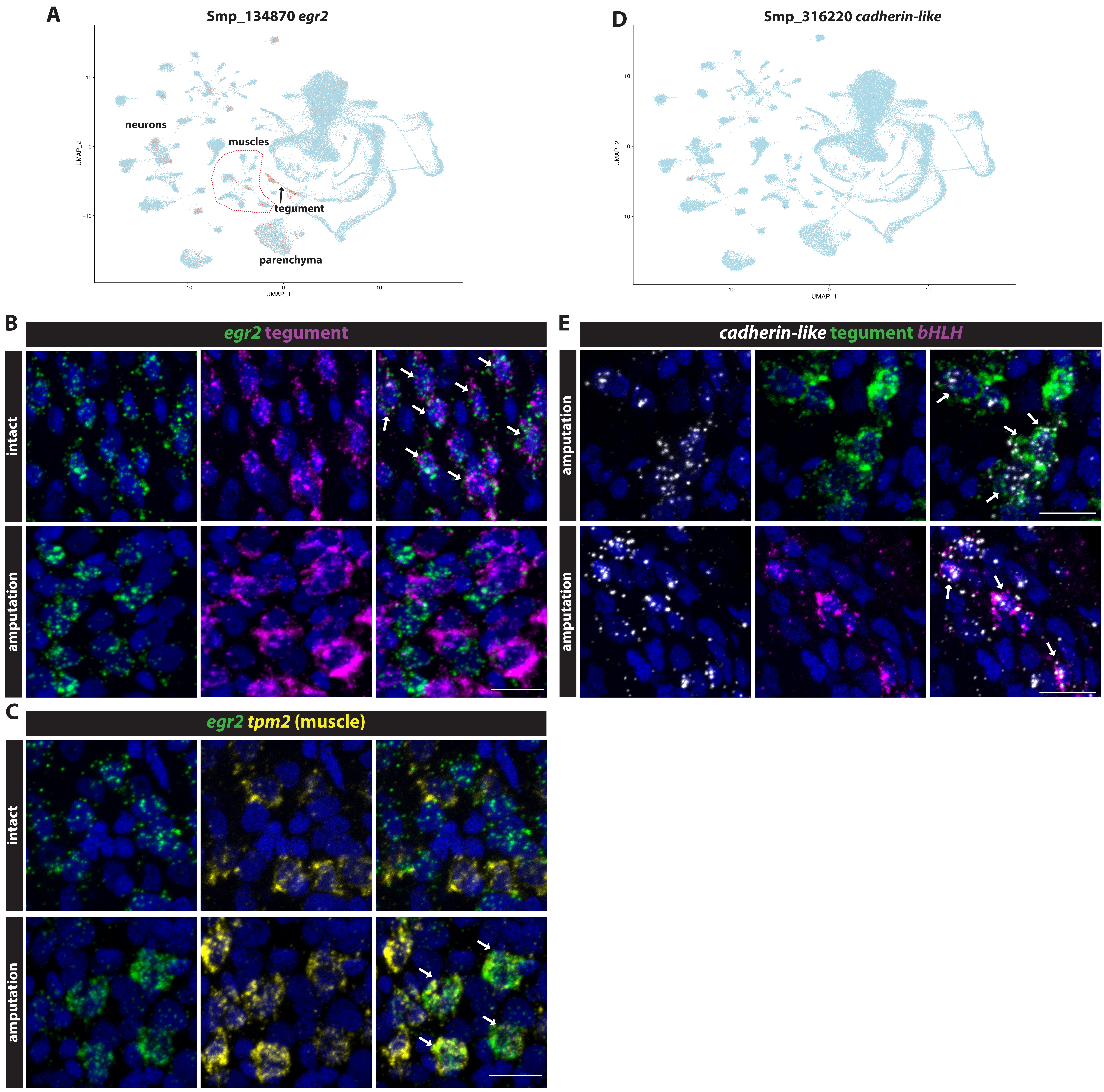

### Figure S5

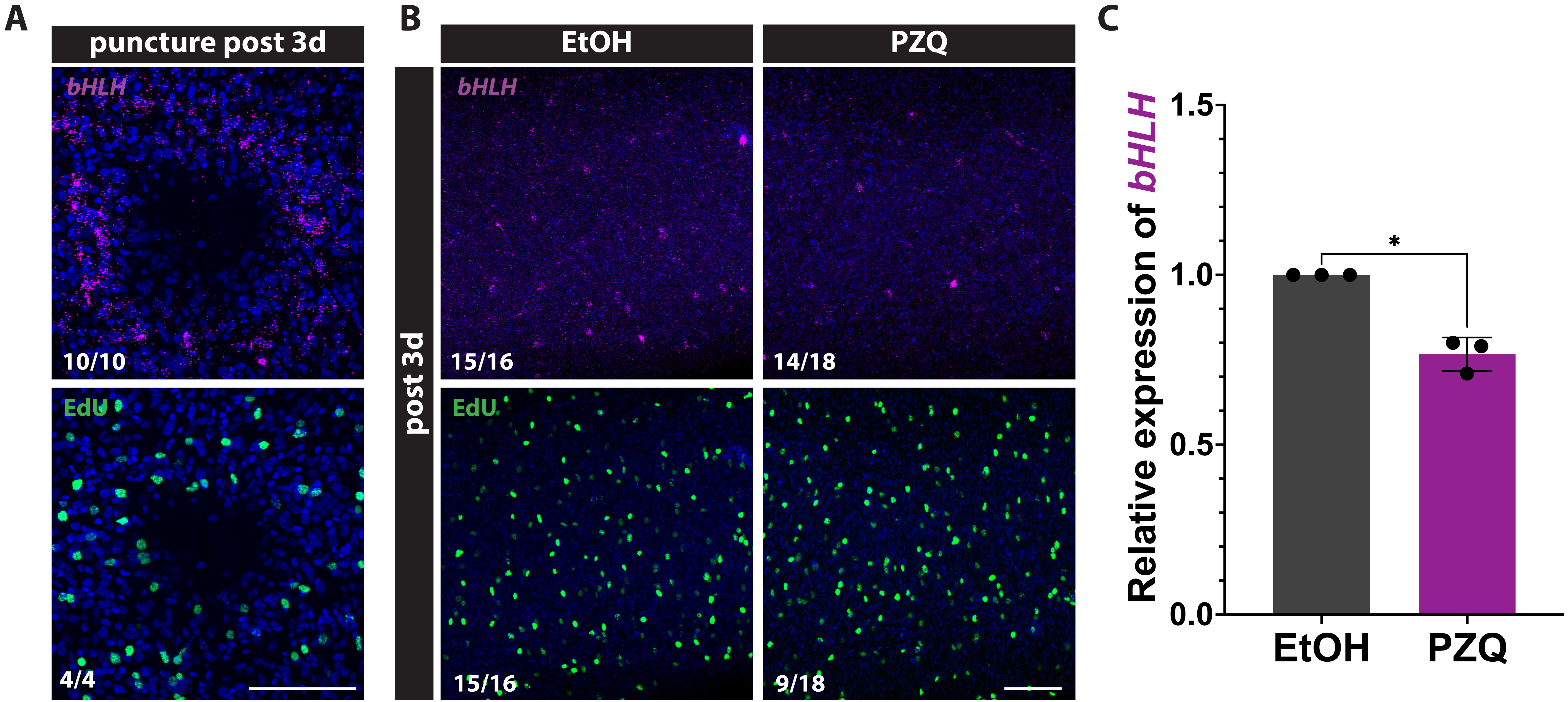

### Figure S6

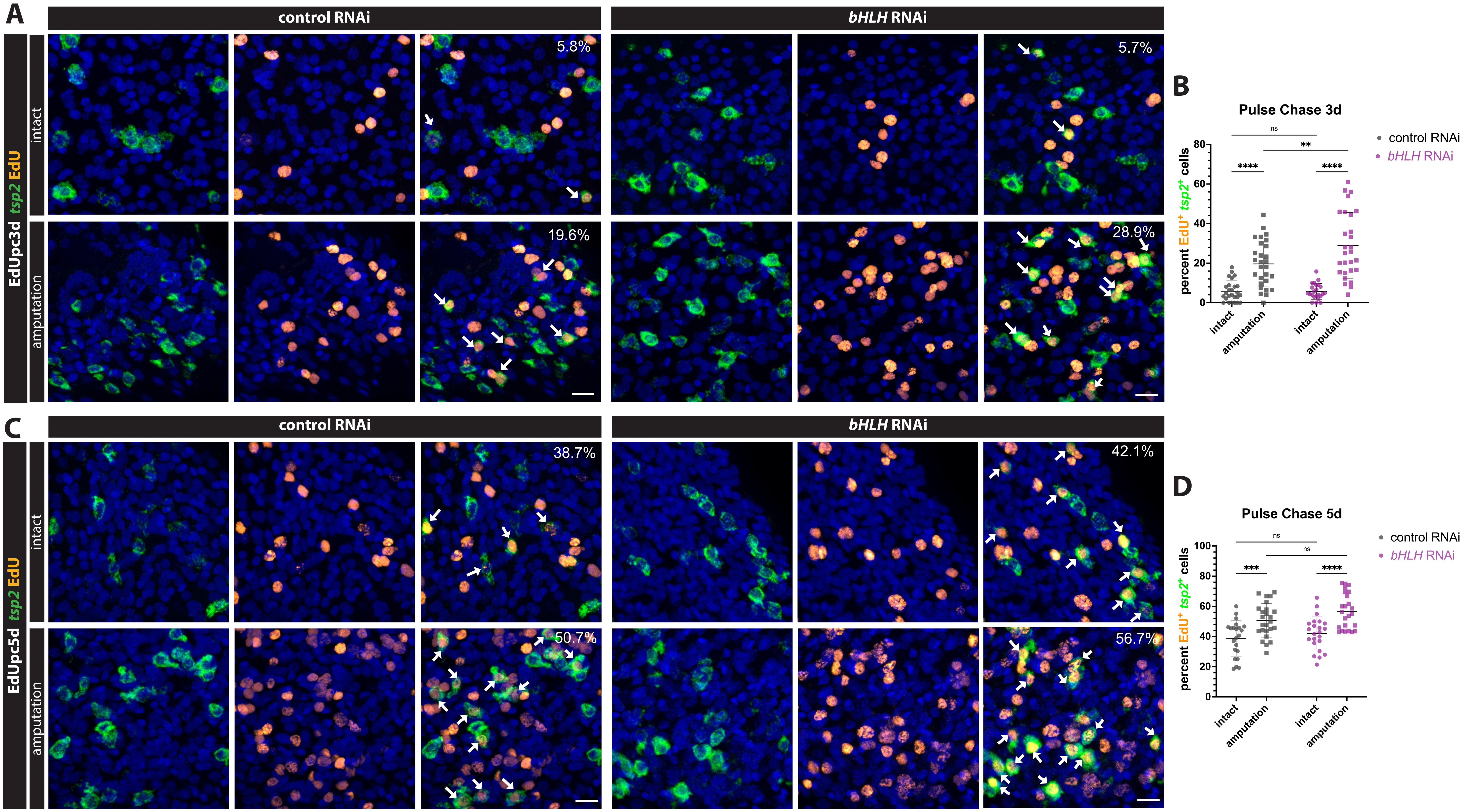

### Figure S7

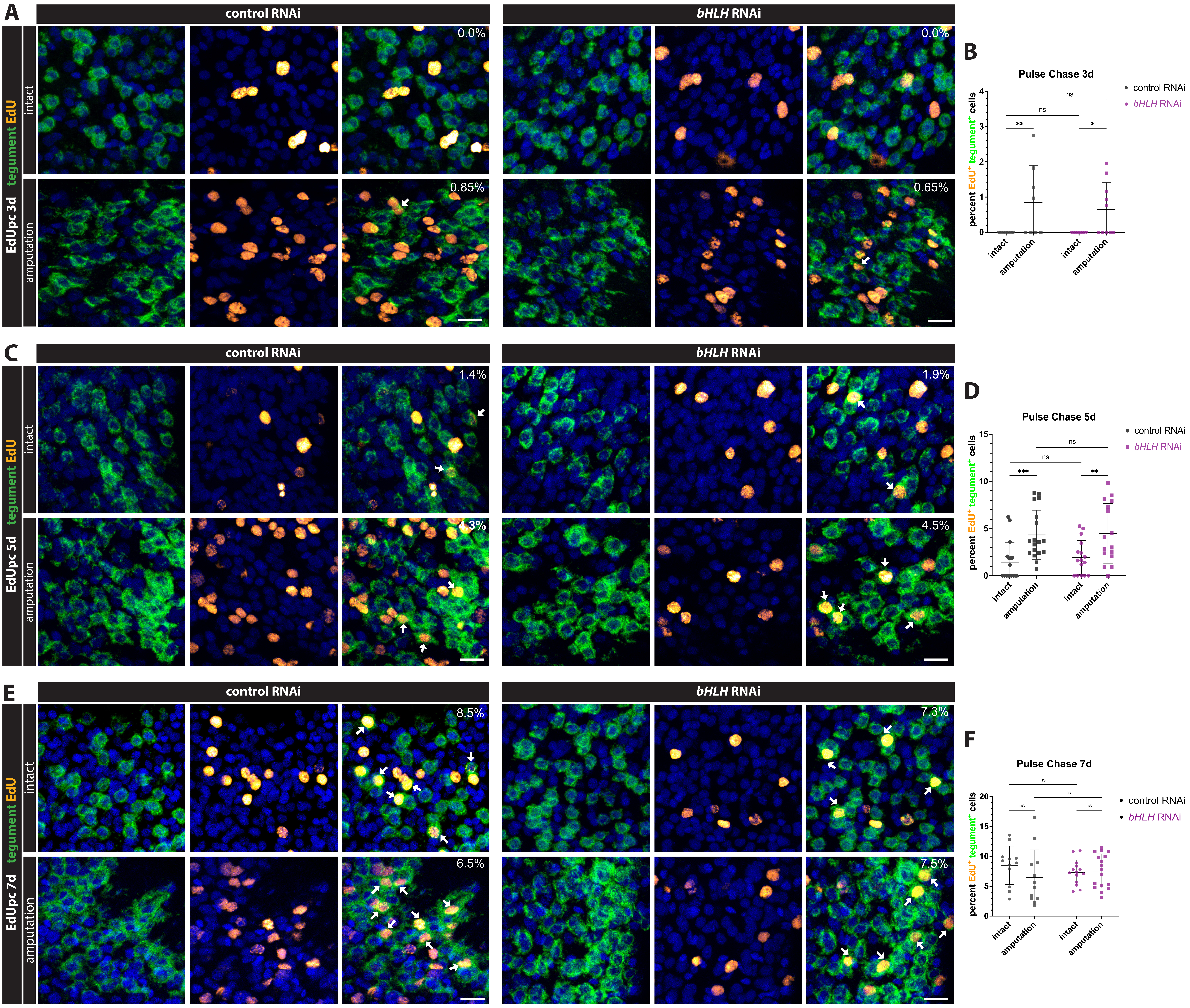

### Figure S9

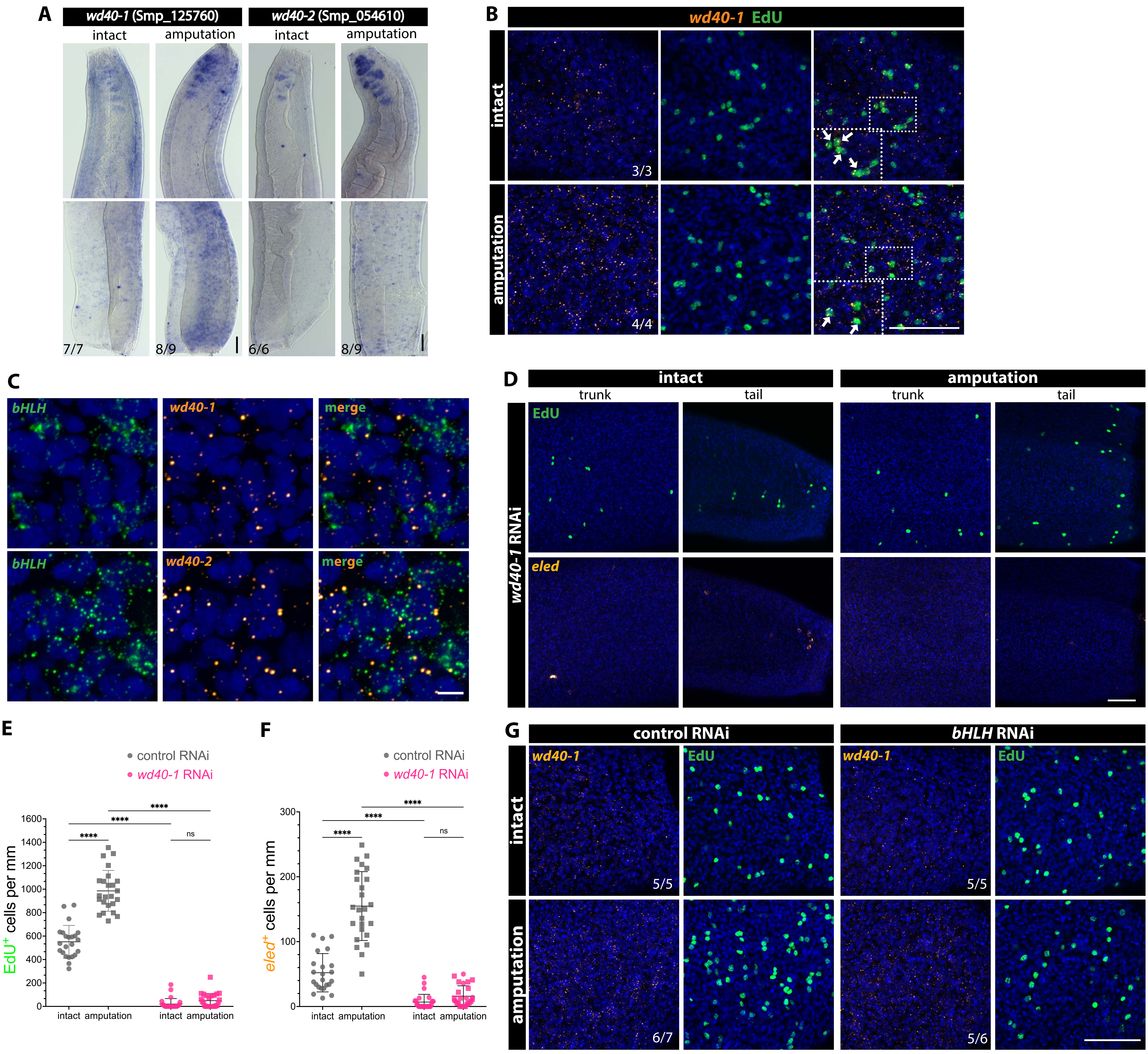

### Figure S10

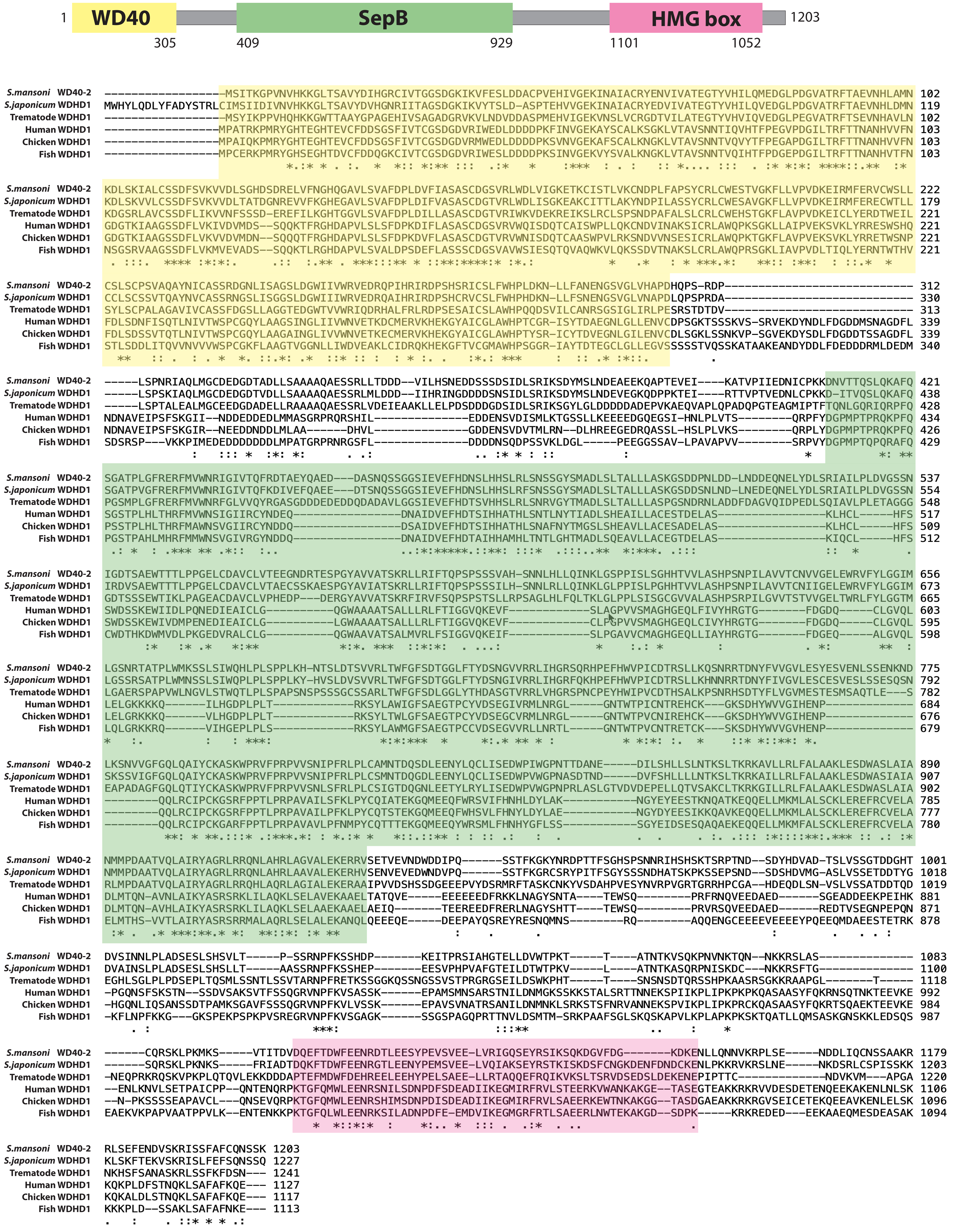

### Figure S11

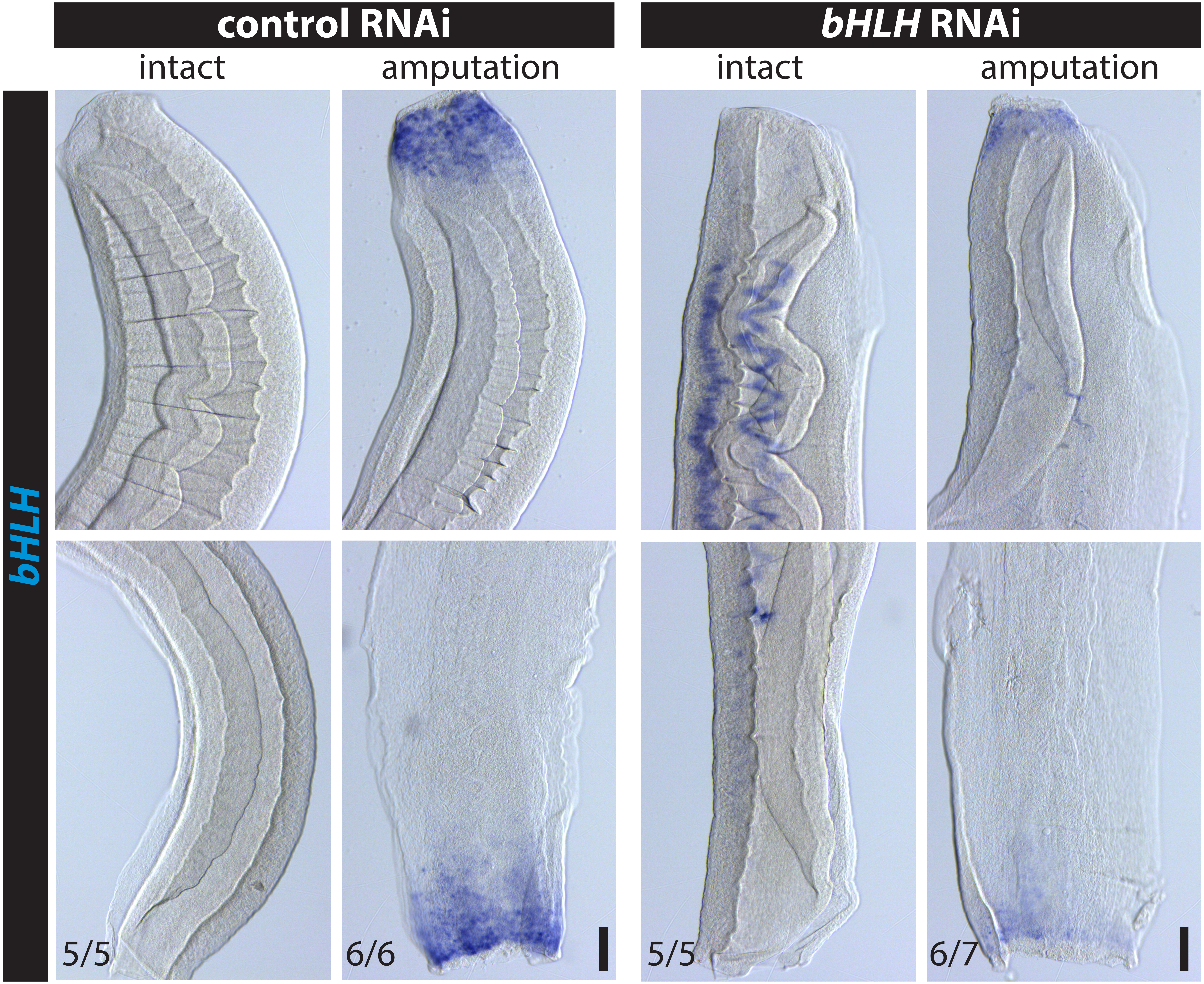
