## Supplementary material for "An injury-responsive *bHLH* is required for regenerative responses following mechanical injury in adult *Schistosoma mansoni*": Figure S8

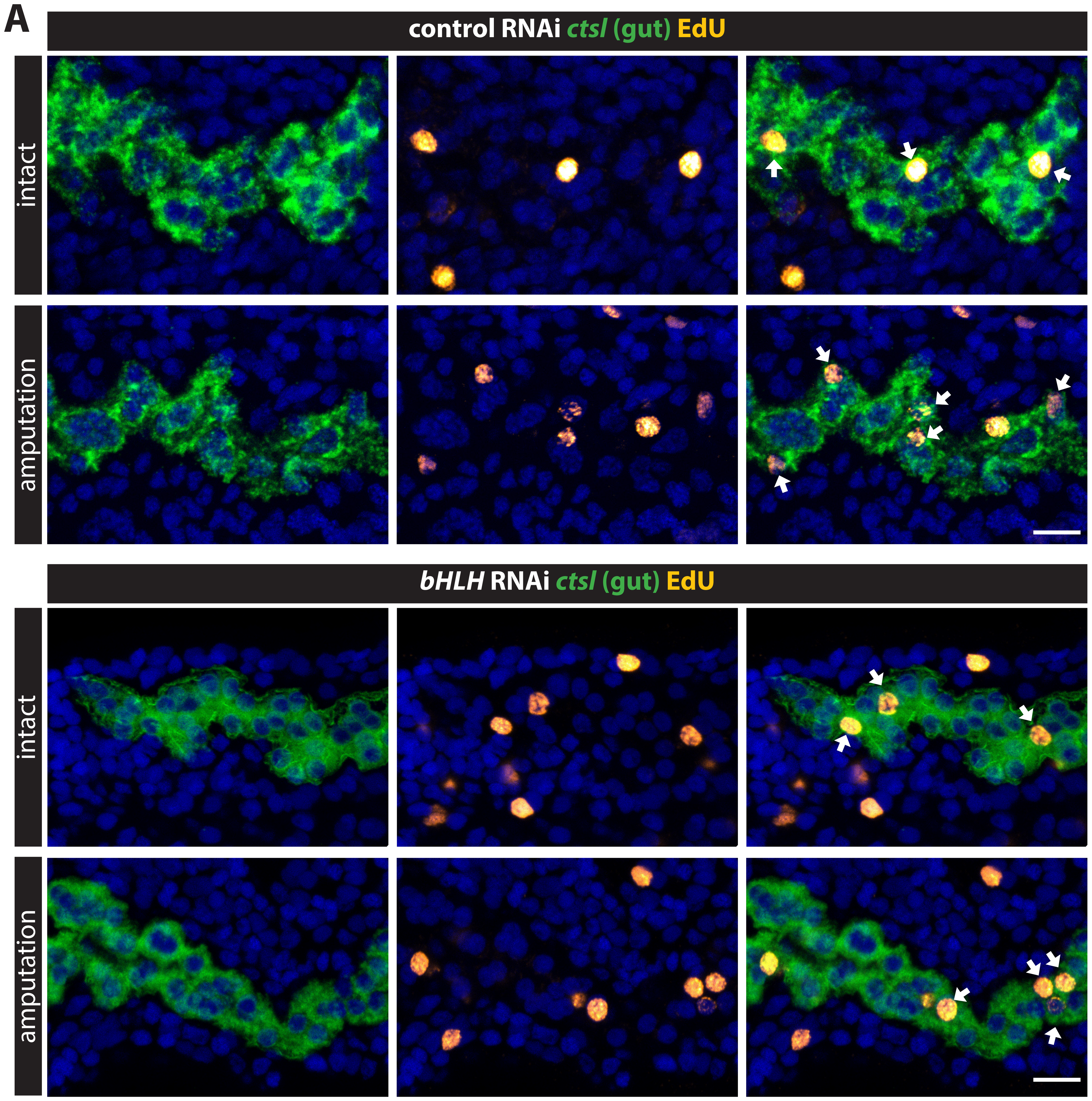


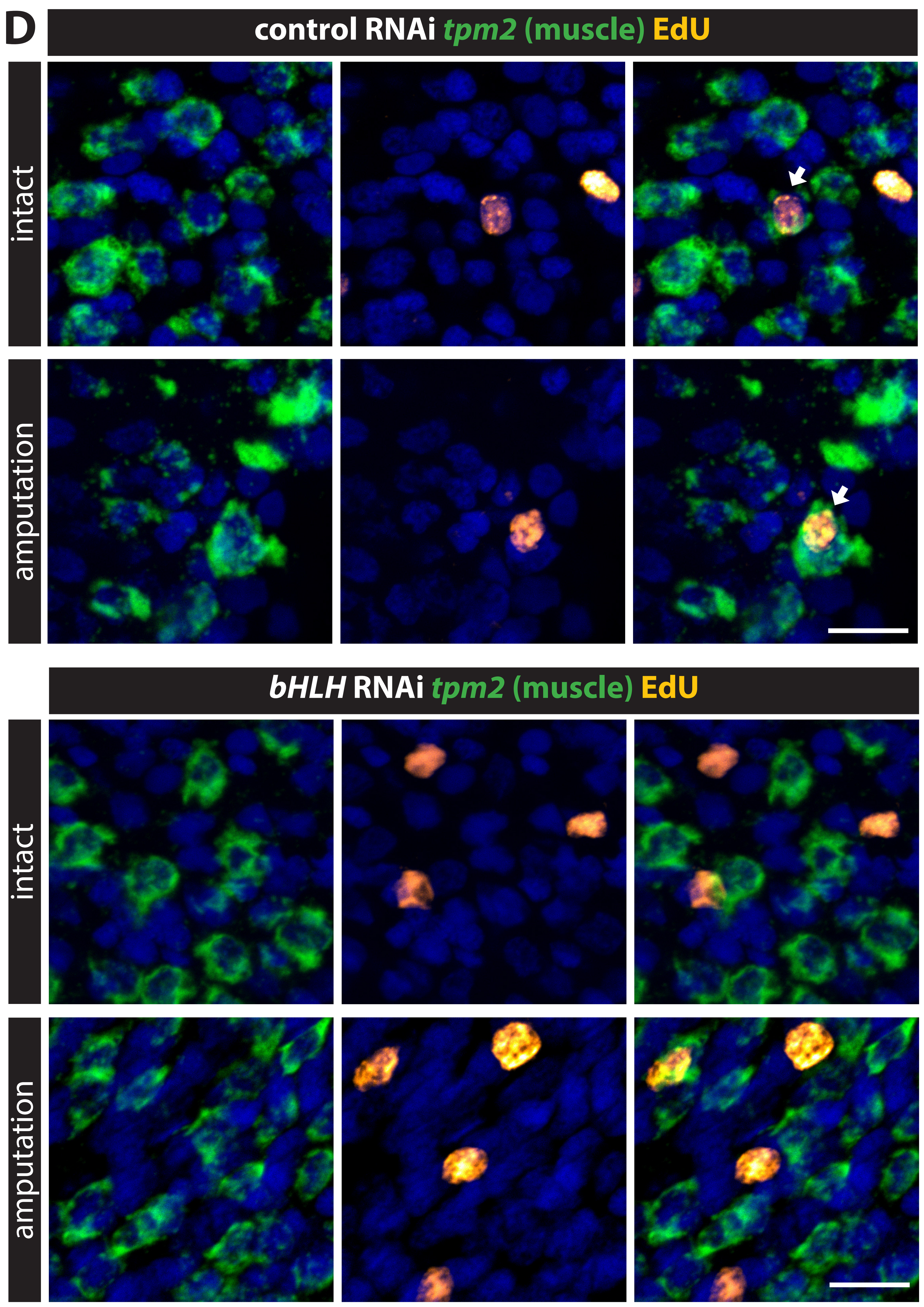


**B**

| | intact<br>EdU⁺intestinal nuclei/DAPI⁺ nuclei (percentage) | amputation<br>EdU⁺intestinal nuclei/DAPI⁺ nuclei (percentage) |
|---|---|---|
| control RNAi | 32/930 (3.44%) n=18 | 57/1006 (5.67%) n=20 |
| *bHLH* RNAi | 55/1114 (4.94%) n=20 | 81/1105 (7.33%) n=24 |

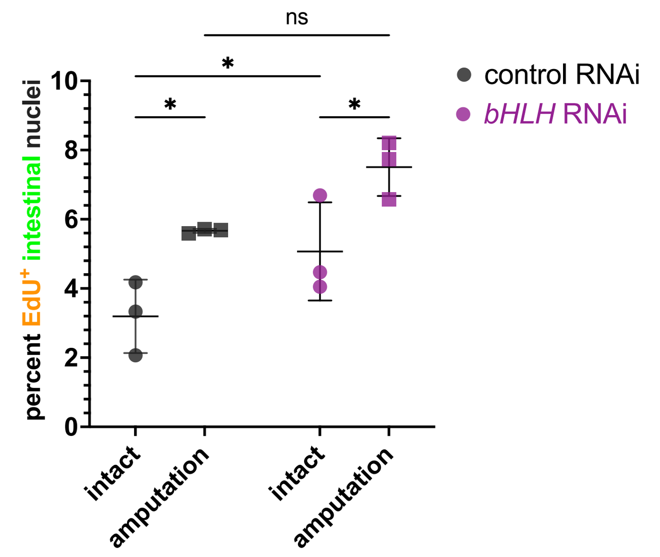


**E**

| | intact<br>EdU⁺muscle cells /muscle cells | amputation<br>EdU⁺muscle cells /muscle cells |
|---|---|---|
| control RNAi | 1/1372 (n=15) | 3/1085 (n=17) |
| *bHLH* RNAi | 1/1184 (n=14) | 0/1039 (n=15) |

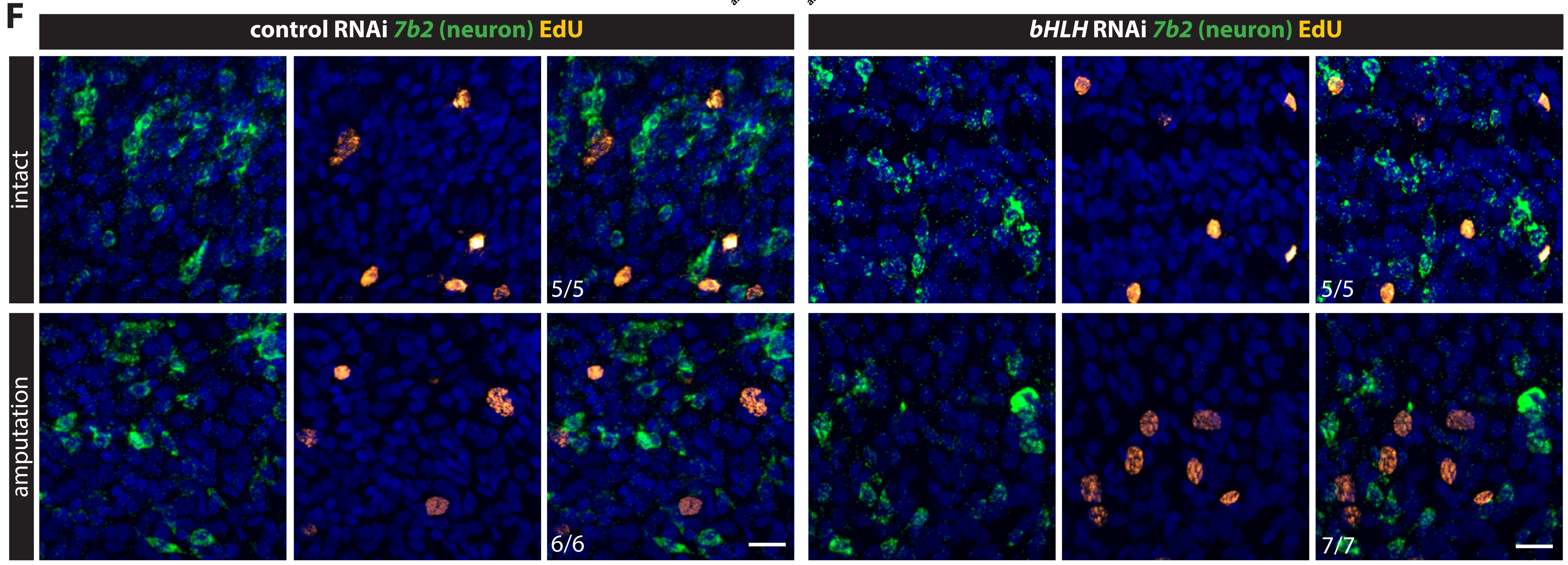
